# Loss of Elp3 impairs the maturation tempo of brain ependymal cells

**DOI:** 10.1101/2025.01.07.631730

**Authors:** S. Tielens, C. Boutin, S. Huysseune, N. Thelen, M. Charlet-Briart, C. Reyskens, L. Oskera, A. Boutsen, S. Mateo-Sanchez, L. Delacroix, P. Labedan, M. Thiry, B. Malgrange, A. Chariot, F. Tissir, C. Creppe, E. Seuntjens, S. Laguesse, L Nguyen

## Abstract

Conditional deletion of *Elp3* in the mouse forebrain leads to microcephaly at birth. In this study, we demonstrate that these mice also develop postnatal hydrocephalus, associated with an enlargement of the brain ventricles. In wild-type mice, ependymal motile cilia are properly aligned to facilitate the circulation of cerebrospinal fluid (CSF) within the ventricles. Our findings reveal that *Elp3* loss induces endoplasmic reticulum (ER) stress and upregulation of ATF4 expression in ependymal cell progenitors, which compromises Notch signaling and accelerates their maturation. This is accompanied by a disruption in the establishment of rotational and translational polarities of the motile cilia of maturing ependymal cells, resulting in disorganized cilia bundles. Collectively, these molecular abnormalities lead to the premature and abnormal development of ependymal cells, culminating in cilia beating dysfunction, impaired CSF clearance, and the development of hydrocephalus.

## Introduction

Ependymal cells are unique, multiciliated cells that form the inner lining of the brain and the spinal cord. In the brain’s ventricular system, these cells play a key role in the central nervous system (CNS) by moving cerebrospinal fluid (CSF) and maintaining brain homeostasis ^1, 2^. At the onset of brain formation, a special layer called the neuroepithelium lines the brain’s ventricles. Later during brain development, the neuroepitelial cells transformed into the radial glial cells (RGCs), which will progressively give rise around birth to the ependymal cells that line the adult mice brain ventricles ^3^. Ependymal cells mature and develop their motile cilia in the first few weeks after birth ^3^. RGCs harbour a primary cilium on their surface that point into the ventricles, which acts like a “sensor,” picking up signals from the outside (e.g. wingless, Wnt, or sonic hedgehog, Shh) and helping regulate a broad array of homeostatic mechanisms ^4–8^. The primary cilium has an axoneme structure, covered by a ciliary membrane connected to the cell membrane. Certain proteins from the centrosome help anchor and stabilize the basal bodies, which are crucial for forming the primary cilium.

The correct positionning of the primary cilium on the surface of RGCs is an important step for the maturation of ependymal cells ^9^. The following stages involve a complex process during which centrioles are multiplied and converted into basal bodies before moving towards the plasma membrane where they dock to form motile cilia ^10–14^. The maturation of RGCs into ependymal cells is controlled by the activity of several transcription factors, including Six3, NFIX, S100b, and FoxJ1 ^15, 16^. In particular, the expression of FoxJ1 is activated by a molecular pathway involving proteins such as Notch, GemC1, McIdas, E2F family members, p73, and c-Myb ^17–22^.

The establishment of a directed flow of CSF at the surface of ependyma relies on the organization of cilia across scales ^23^. Two types of polarity have been described in ependymal cells^1, 9^. At the cellular scale, translational polarity refers to the clustering of basal bodies into an off-center patch located on one side of the apical surface ^9, 24^, while rotational polarity refers to the coordinated orientation of basal bodies ^9, 25–28^. At the tissue level, both polarities are coordinated, with neighbouring cells having basal bodies patch localisation and basal body orientations in the same direction ^23^.

Failure of ependymal cells differentiation and function often leads to hydrocephalus, a condition where CSF builds up in the brain’s ventricles ^29^. This is a common central nervous system disorder, affecting about 4.65 in every 10,000 births ^30^. Studies in mice have shown a strong link between ciliopathies and hydrocephalus ^26, 31, 32^.

Elp3 is the enzymatic subunit of the Elongator complex ^33^, which is crucial for controlling key cellular functions during brain development such as neurogenesis and neuronal migration ^34–38^. At the molecular level, Elongator promotes wobble uridine modification of selected tRNA species, directly influencing RNA translation efficiency ^39–43^. In previous research, we showed that conditionally deleting Elp3 in the forebrain triggers the PERK/ATF4 pathway of the unfolded protein response (UPR) as a result of endoplasmic reticulum (ER) stress, leading to microcephaly ^34^. In this study, we demonstrate that loss of Elp3 in maturing ependymal cells causes postnatal hydrocephalus. This condition stems from an early defect in primary ciliogenesis in RGCs, followed by improper establishment of rotational and translational polarities in ependymal cells. At the molecular level, the absence of Elp3 results in defective maturation of Notch receptors, disrupting the timing of ependymal cell maturation.

## Results

### Depletion of Elp3 in the mouse forebrain results in postnatal hydrocephalus

We used a previously described conditional knock-out mouse model to delete Elp3 in forebrain FoxG1^+^ progenitors. These mice (referred to as Elp3cKO) exhibited severe microcephaly at birth ^34^. These mice were growth retarded, and most did not survive past weaning (**Figures S1a, S1c**). Notably, we noticed a striking enlargement of the lateral ventricles on brain sections from postnatal day (P) 21 Elp3cKO animals, which signs hydrocephalus (**Figure 1a**). No sign of enlarged ventricles was observed at birth (**Figure S1d**), indicating that the deletion of Elp3 in the developing forebrain later leads to postnatal development of hydrocephalus. Loss of Elp3 in the brain lateral wall after birth was confirmed by immunolabeling (**Figures 1b-c**), RT-qPCR, western blot and mRNA labeling (**Figures S1b, S1e, S1f**). Analysis of single cell RNA sequence dataset from P0 mouse brain confirmed the expression of Elongator subunits in ependymal cells and RGCs, their progenitors (**Figure 1d**) ^44^. Hydrocephalus often results from poor circulation of CSF due to impaired ependymal cell function ^45^. We thus assessed ependymal wall generated CSF flow by monitoring the movement of fluorescent beads moving across the surface of dissected lateral walls from P16 WT and Elp3cKO mice (**Figures 1e-1g**). As previously described ^23^, we observed two streams of beads converging rostro-ventrally at the surface of WT lateral walls. In contrast, beads swirled along various uncoordinated paths across Elp3cKO lateral walls (**Supplemental Movies 1-2**) demonstrating that flow establishment is impaired at the surface of Elp3cKO ependyma. We further examined P16 brain ventricular walls by scanning electron microscopy to analyse cilia in ependymal cells. The cilia bundles were consistently interspaced in the WT lateral walls. In strike contrast, the Elp3cKO ependymal cells show disorganized cilia tufts that were spaced irregularly across the lateral wall surface. Additionally, individual cilia appeared intermingled within cilia bundles (**Figures 1h-1i**). In some patches, the tips of several cilia were stuck together, forming sheet-like structures (**Figure 1i**, arrow in close up). The length of cilia per bundle were not affected in Elp3cKO as compared to WT (**Figures 1j**). Together, these data show that the lack of Elp3 expression leads to disorganized cilia tufts at the lateral wall surface, which impairs CSF flow circulation generating postnatal hydrocephalus in mice.

**Figure 1.**
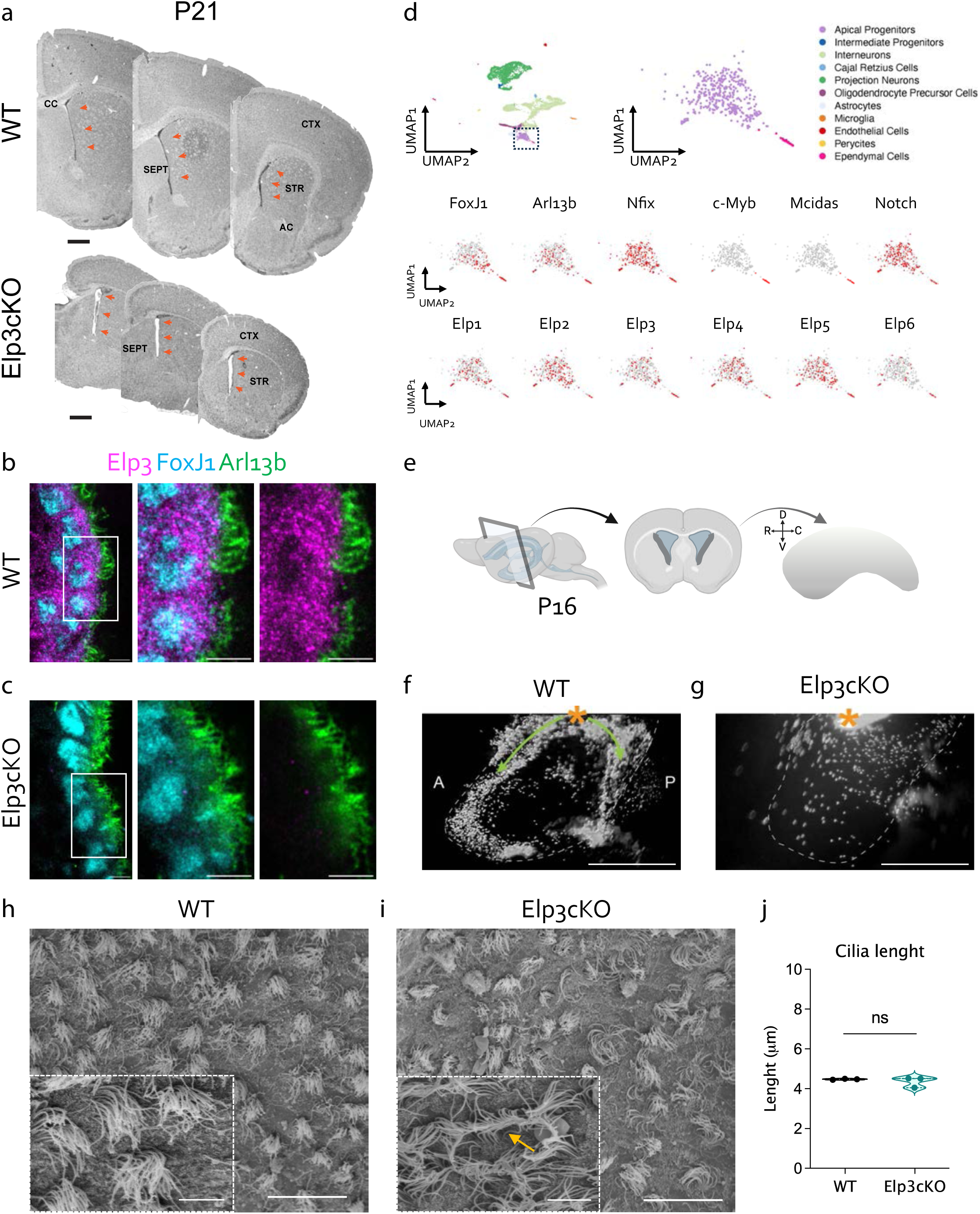
Deletion of Elp3 in the forebrain causes postnatal hydrocephalus in mice. **a**, Consecutive coronal sections along the rostrocaudal axis of P21 WT and Elp3cKO mouse brains showing enlargement of Elp3cKO brain lateral ventricles (luminal space pointed by orange arrows). Scale bars, 500 µm. **b,c,** Immunolabeling of P7 lateral wall from WT or Elp3cKO mice showing expression of Elp3 (magenta), Foxj1 (blue), and Arl13b (green). Scale bars, 5 µm. **d**, bioinformatic analysis of published single cell data showing the connected clusters of apical progenitors and ependymal cells in microdissected lateral wall form P0 mice ^44^. The distribution of a selection of ependymal cell genes together with Elongator subunit genes are shown. **e,** schematic representation of dissected lateral wall from P16 mouse brain. **f,g,** Representative images illustrating displacement of floating fluorescent beads at the surface of WT (**f**) and Elp3cKO (**g**) immerged brain lateral walls. Dashed lines delineate surface of brain lateral wall explant, orange star corresponds to the initial bead deposit. Green arrows represent the physiological CSF streams, which are disorganized in Elp3cKO. Scale bars, 500µm. **h,i,** Scanning electron microscopy micrographs at the surface of WT (**h**) and Elp3cKO (**i**) brain lateral wall. Close ups show ciliary patch organization, which is lost in Elp3cKO. Scale bars, 20µm on the main microph and 5µm in the close up. **j,** Quantification of cilia length in WT and Elp3cKO P16 lateral wall. Violin plots show median ± quartiles from three biological replicates. p-values were determined by Mann-Whitney test. CTX: cortex; CC: corpus collosum; SEPT, septum; A: anterior, P: posterior.; LW, lateral wall, MW, medial wall, AC, anterior commissure;. Scale bars: A: 500 µm; B: 20 µm; C: 5 µm.

### Loss of Elp3 perturbs the establishment of cilia polarities in ependymal cells

Brain ependymal cells organize their motile cilia to generate unidirectional CSF flow. To further understand the underlying mechanisms of hydrocephaly in Elp3cKO, we analyzed cilia polarities which are reflected by basal bodies (BB) organization ^1^. We performed immunolabeling on the dorsoventral region of the lateral wall using antibodies against γ-tubulin that marks the basal foot, FGFR1 Oncogene Partner (FOP) which stain the centriolar wall of the BB ^46^, and ZO1 that delinates ependymal cell borders. This triple staining allow the concomitant analysis of translational and rotational polarity at cell and tissue levels ^23, 24, 47^.

First, we analyzed rotational polarity. At cell scale, FOP and γ-tubulin allowed us to draw for each BB a vector (Vcil) which reflects the beating direction of the cilia (arrows in **Figures 2a-d**). In addition, we defined the vector of patch orientation (VpatchO), which represents the mean direction of all individual Vcil in a given cell (**Figures 2c, 2d**). At P16, BBs within WT ependymal cells were aligned in parallel rows, regularly spaced, and all Vcil pointed in a similar direction which resulted in a low circular standard deviation (CSD) (**Figure 2a, 2c, 2e**). In contrast, Elp3cKO ependymal cilia were not uniformly aligned, showed uncoordinated Vcil orientation (**Figures 2b, 2d**). This was reflected in significantly higher CSD of Vcil vectors compared to WT mice (**Figures 2e**). VpatchO was determined in cells with a CSD below 40° ^23^ and was only reported for 77.8% of Elp3cKO ependymal cells compared to 99.8% of WT ependymal cells (**Figure 2e**).

**Figure 2.**
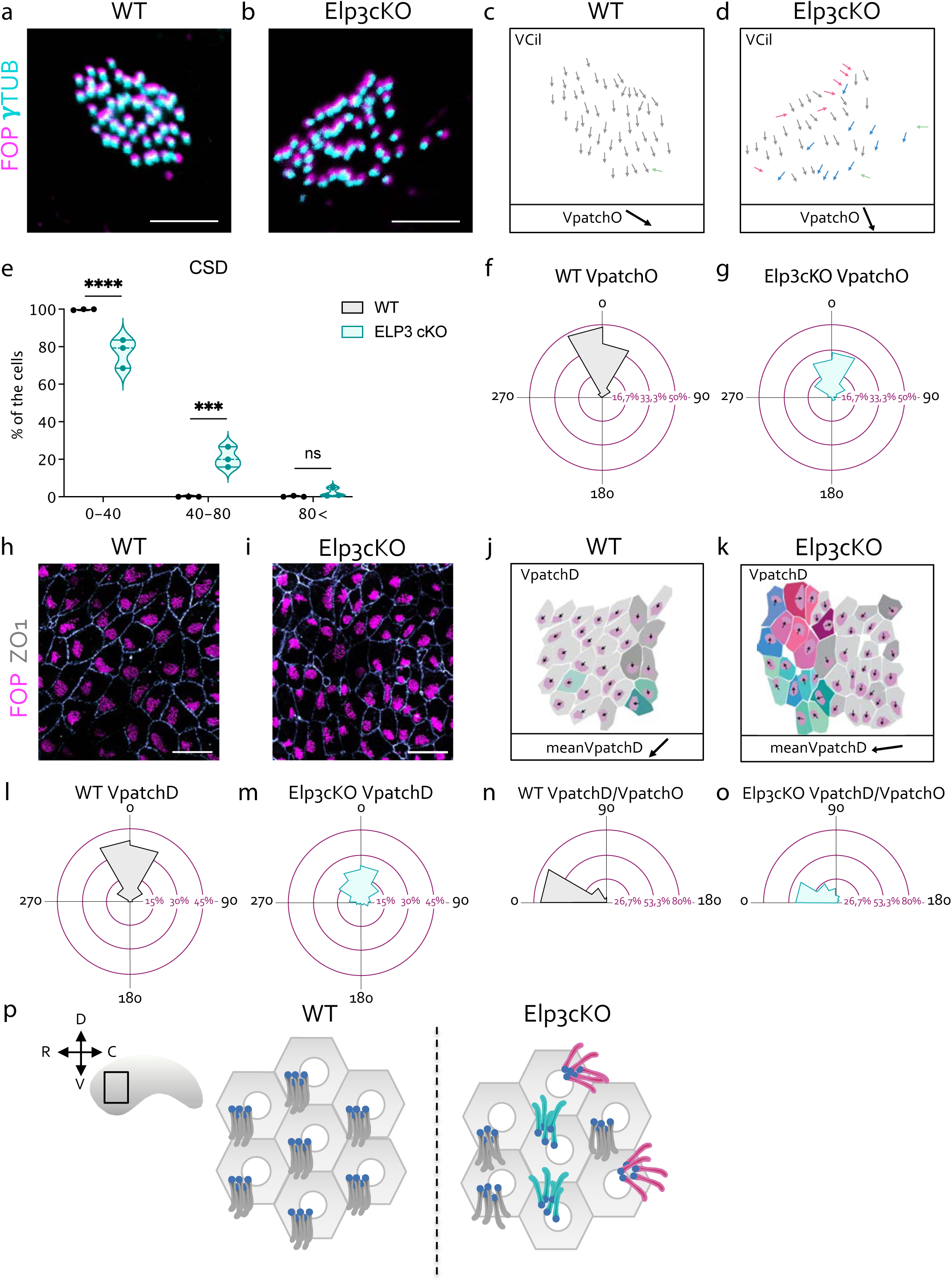
Loss of Elp3 perturbs motile cilia rotational and translational polarities. **a,b,** Confocal images of motile cilia patches labelled with FOP (magenta) and γ-tubulin (green) on WT (**a**) and Elp3cKO (**b**) on the dorsoventral region of the brain lateral wall of P16 mice. Scale bars, 3µm. **c,d,** Analysis of the rotational polarity of motile cilia patches. Representative ciliary vector (VpatchO) represents the average polarity direction of independant motile cilia within a cellular patch in a WT (**c**) and a Elp3cKO (**d**) maturating ependymal cell. Pseudocolors representing individual motile ciliary vectors. **e**, Percentage of ependymal cells according to the ciliary vector standard deviation (CSD) in WT and Elp3cKO P16 lateral wall. Violin plots show median ± quartiles from three biological replicates. P-values were determined by two-way ANOVA with Bonferroni’s multiple comparisons test \*\*\*\**P*<0.0001 and \*\*\**P*<0.001. **f,g,** Circular dispersion plots of VpatchO around the mean in WT (446 cells) (**f**) and Elp3 cKO (440 cells) (**g**) from three biological replicates. p-values were determined by Watson’s U^2^ test=3.683; ***p<0.001. **h,i,** Confocal images of motile cilia patches labelled with FOP (magenta) and ZO1 (light grey) on WT (**h**) and Elp3cKO (**i**) lateral wall. Scale bars, 20µm. **j,k,** Analysis of the translational polarity of motile cila patches amongst ependymal cells. VpatchD designates the average translational vector of multipe neighbouring ependymal cells, which indicates the displacement direction of BB patch according to the ependymal cell geometrical center. The quantification is based on immunolabeling of WT (**j**) and Elp3cKO (**k**) lateral wall surface area. Pseudocolors correspond to individual BB patch displacement directions. **l, m,** Representation of the circular dispersion of individual VpatchD around the mean in WT (446 cells) and Elp3cKO (466 cells); brain lateral wall area from six and eight biological replicates, respectively. p values were determined by Watson U^2^ test=1.151, *** p<0.001. **n,o,** Circular dispersion of the correlation between VpatchD (translational polarity) and VpatchO (rotational polarity) angles in WT (445 cells) and Elp3 cKO (355 cells); from three biological replicates. p values were determined by Watson U^2^ test=1.714, ***p<0.001. **p,** Schematic representation of the translational and rotational polarity defects defects upon Elp3 loss compared to WT ependymal cells. FGFR1 Oncogene Partner (FOP).

At the tissue level, neighboring ependymal cells present coordinated BB patch orientation (tissue-level rotational polarity) ^27, 28, 48, 49^. To quantify this, we compared the VpatchO of all neighboring ependymal cells to the meanVpatchO of the analyzed field. In WT, the low circular dispersion of VpatchOs around the mean indicated a coordinated intercellular orientation (**Figure 2f**). In contrast, in Elp3cKO lateral walls, the coordination of patch orientation was lost, wich resulted in a significantly larger distribution of VpatchO vectors (**Figure 2g**). Overall, these results show that loss of Elp3 alter both intra- and intercellular orientation of ependymal cilia.

Next, we examined whether loss of Elp3 impacts the establishment of the translational polarity using FOP and ZO1 staining (**Figures 2h-2i**) as decribed previously ^23^. A vector “VpatchD” (arrows in **Figures 2j-2k**) indicating the displacement of the BB patch, was drawn from the geometrical center of each cell’s apical surface to the center of the BB patch. In WT, neighboring ependymal cells showed similar directions of BB patch displacement, reflected by the alignment of individual VpatchDs, all pointing to the same direction (color-coded cells in **Figure 2j**). In contrast, Elp3cKO lateral walls displayed altered BB patch organization and clusters of cells coordinated in opposite directions were observed (**Figure 2k**, color-coded cells). This resulted in a much broader circular dispersion of VpatchD around the mean in Elp3cKO mice compared to WT mice (**Figures 2l-2m**). This result indicated that the coordination of translational polarity is impaired upon loss of Elp3.

Finally, we assessed wether the intracellular coupling of BB patch orientation and displacement observed in WT ^23^ was preserved in Elp3cKO. The angles between VpatchD and VpatchO were measured in WT and Elp3cKO cells. In WT ependymal tissue, VpatchD/VpatchO angles were below 45° in 70% of ependymal cells (**Figure 2n**). In contrast, in Elp3cKO ependymal cells, the angles were significantly larger (up to 180°), with a 30% reduction in the number of cells exhibiting angles below 45° (**Figure 2o**). This demonstrated an uncoupling between BB patch orientation and displacement in Elp3cKO deficient cells. Altogether, our data demonstrate that the lack of Elp3 results in a disorganization across multiple scales of ependymal cell polarities (summarized in **Figure 2p**).

### Loss of Elp3 impairs tissular distribution of Planar Cell Polarization (PCP) proteins across the brain lateral wall

At the molecular level, establisment of ependymal polarity relies on coordinated distribution of Planar cell polarity (PCP) proteins which is integrated by ependymal cells to organize their basal bodies ^23, 24, 50^. To analyse whether Elp3 depletion impacts PCP, we analyzed the expression level of key core PCP genes (*Ankrd6*, *Celsr1*, *Celsr2*, *Celsr3*, *Fzd3*, *Fzd6*, and *Prickle1*) and found no differences between P16 WT and Elp3cKO brain lateral walls (**Figure S2a**). Next, we analysed the distribution of Vangl2 in WT and Elp3cKO lateral walls (**Figures S2b-S2c**). In both conditions, Vangl2 labeling was detected in a small fraction of the cell perimeter that mainly localized on the opposite side of the ciliary patch. To assess the coordination of Vangl2 distribution across the tissue, we drew a vector (Vvangl2) from the center of the ciliary patch to the center of the Vangl2 staining (**Figures S2d-S2e**). In WT mice, we observed a coordinated localization of Vangl2 across the tissue, reflected by low circular dispersion of Vangl2 vectors (**Figure S2f**). In contrast, this coordination was altered in Elp3cKO mice (**Figure S2g**). Coordination of Vangl2 distribution was preserved in small clusters of cells. However, adjacent clusters showed opposite Vangl2 protein localization. This organization echoes a defect in translational polarity as described above. These findings show that Elp3 depletion does not affect neither PCP gene expression nor protein asymmetric localization in individual cells, but rather disrupts the coordination of protein distribution at the tissue level.

### Loss of Elp3 affects primary ciliogenesis in radial glial cells

Ependymal cells originate from the differenciation of RGCs, which are organized in a sheet-like structure ^3, 24, 51^. RGCs possess a non-motile single primary cilium that extends into ventricles. Coordinated asymmetrical positioning of the primary cilium at the apical surface of RGCs is essential for the proper planar polarization of the future ependymal cells ^9, 24, 52^.

To investigate the positioning of the primary cilium in RGCs, we performed co-immunolabeling for ZO1 (to delineate cell borders) and γ-tubulin (marking the BBs of primary cilia) on P0.5 WT and Elp3cKO lateral walls (**Figures 3a-3b**). We assessed the position of the primary cilium in respect the cell center to define a polarity vector (VD) and measured its circular deviation around the mean (meanVD) of neighbouring RGCs (arrows in **Figures 3c-3d**). Most vectors in WT pointed in a comparable direction, with the angular deviation around the mean ranging from −45° to +45° (**Figure 3e**) ^9^. In contrast, the angular deviation in Elp3cKO mice had a significantly broader dispersion (**Figure 3f**). Notably, no RGC clusters showed coordinated polarization in the lateral wall of P0 Elp3cKO mice (**Figure 3d**). This shows that loss of Elp3 impairs the coordination of basal body positioning between neighboring RGCs.

**Figure 3.**
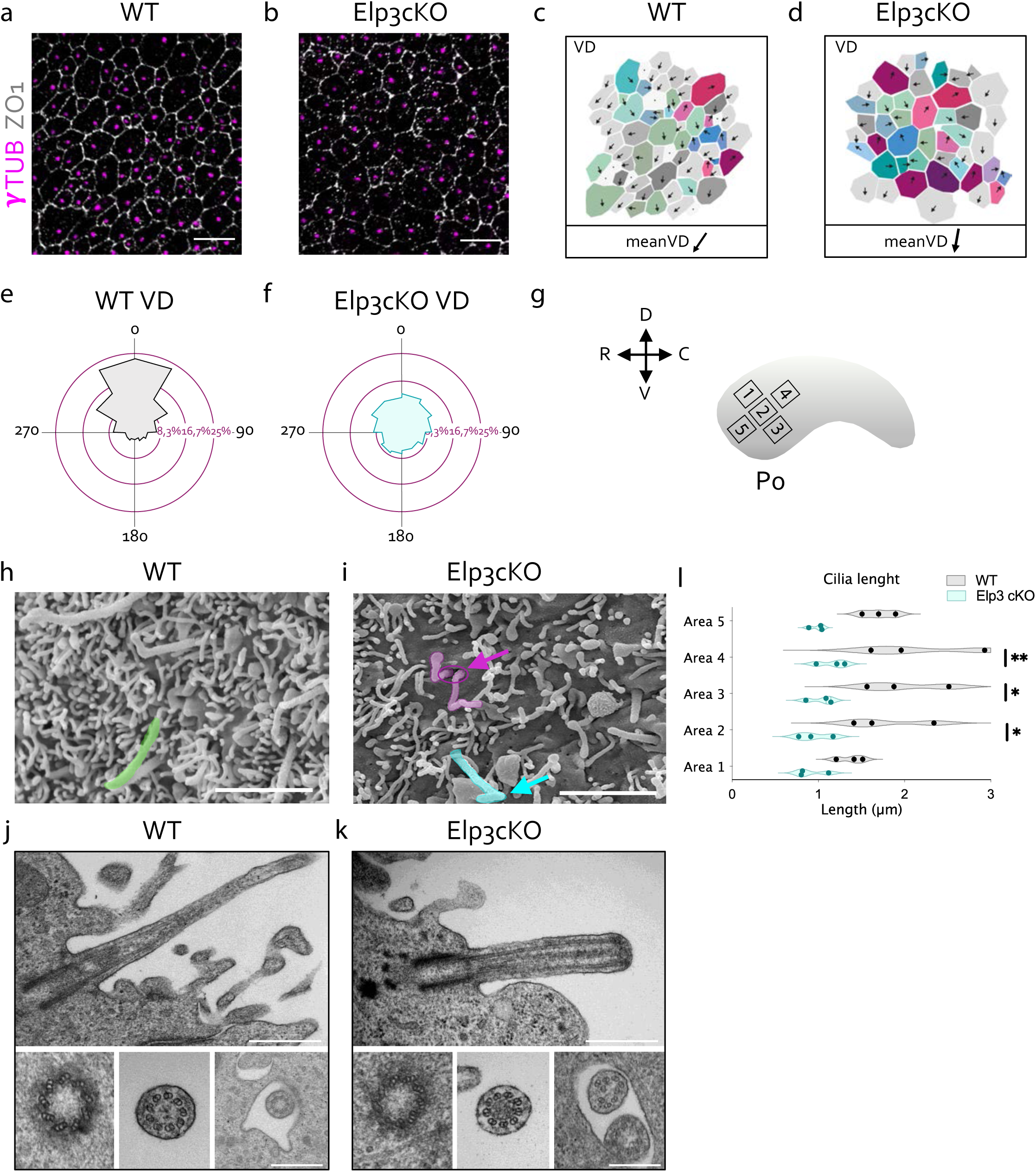
Loss of Elp3 affects primary ciliogenesis in radial glial cells. **a,b,** Immunolableings of radial glial cells (RGCs) showing the enchorage of the RGC primary cilium expressing γ-tubulin (magenta) and the RGC perimeter expressing ZO1 (light grey) in the brain lateral wall of P0.5 WT (**a**) and Elp3cKO (**b**) mice. Scale bars, 10µm. **c,d,** Pseudocolored arrows representing BB displacement vectors (Vcil). The MeanVD is the mean of the individual Vcils. **e, f,** Representation of the Vcils at the tissue level represented as angular dispersion around the MeanVD. WT (2068 cells), Elp3 cKO (2103 cells) from the brain lateral wall of six biological replicates. p values were determined by Watson U^2^ test=9.347, *** p<0.001. **g,** Scheme of the brain lateral wall with squared areas analyzed by electron microscopy (EM). **h,i,** Scanning EM micrographs of P0.5 WT (**h**) and Elp3cKO (**i**) lateral wall showing primary cilia, colored arrows point to defective cilia (green, normal cilium; blue, abnormal kinked cilium; purple, two abnormal kinked cilia sharing one cilium pocket) Scale bars, 2µm. **j,k,** Transmission EM of WT (**j**) and Elp3 cKO (**k**) P0.5 lateral wall showing the ultrastructure of the primary cilium. Close up represent from left to right: basal body, axoneme, and ciliary pocket. Scale bars are 1µm in the micrograph and 200nm in corresponding close ups. **l,** Averaged primary cilium length within squared surface (see also g) of P0.5 for WT and Elp3cKO lateral walls. Violin plots show median ± quartiles from three biological replicates. p-values were determined by two-way ANOVA with Bonferroni’s multiple comparisons test \*\**P*<0.01 and \**P*<0.05.

We then analysed RGC primary cilia by scanning and transmission electron microscopy in five adjacent areas of the dorsoventral region of the P0.5 lateral wall (**Figure 3g**). This analysis revealed severe morphological abnormalities in the primary cilia of Elp3cKO mice, including shorter, kinked and supernumerary (two per ciliary pocket) primary cilia (arrows in **Figures 3h-3i**, right close up in **Figure 3k**). The loss of Elp3 resulted in shorter primary cilia (**Figure 3l**). Transmission electron microscopy confirmed the reduced cilia length in Elp3cKO mice, but their overall ultrastructure was mostly conserved. Indeed, both WT and Elp3cKO cilia contained nine triplets of tubulin at the basal body level and nine doublets in the axoneme (**Figures 3j-3k**). Notably, some Elp3cKO cilia displayed an unknown darker substance within the center of the axoneme (central close up in **Figure 3k**). In addition to primary cilia defects, we also observed a significant reduction in microvilli across the five different areas of the lateral wall surface in Elp3cKO P0.5 pups (**Figures S3a-S3d**). These findings show that Elp3 is essential for proper morphogenesis of the primary RGC cilium.

### Elp3 depletion impairs primary ciliogenesis by interfering with Notch signaling

Next, we aimed to uncover the mechanism by which the loss of Elp3 perturbs primary ciliogenesis. The primary cilium integrity relies on stabilization of its tubulin cytoskeleton, which is associated with some post-translational modifications of α-tubulin, including acetylation. The expression level of acetylated α-tubulin in primary cilium microtubules, which is disrupted in cortical projection neurons lacking Elp3 ^38, 53^, was comparable between genotypes (**Figures S4a-S4c**). Previous studies, have emphasized the key role of Elongator in modifying wobble uridines (U34) at selected cytoplasmic tRNAs, which enhances translation efficiency and accuracy ^34, 54–56^. Specifically, a loss of Elongator activity in cortical progenitors impairs some tRNA modifications, wich causes codon-specific translational pausing. This in turn induces endoplasmic reticulum (ER) stress, leading to the activation of the Unfolded Protein Response (UPR) and deregulation of the neurogenic output in cortical progenitors ^34, 54^. We thus tested whether Elp3 deletion would also triggers ER stress and UPR activation in maturating ependymal cells from the brain lateral wall of P0.5 mice. The analysis of maturating ependymal cells in brain lateral wall revealed highly inflated ER with loss of associated ribosomes in Elp3cKO, which are hallmarks of ER stress (**Figures 4a-4b**). Moreover, ATF4, a key transcription factor - whose expression is activated upon ER stress in cortical progenitors ^34^ - was enriched in protein extracts from the lateral walls of newborn Elp3cKO brains (**Figure 4c**) together with its direct target, the ER stress-inducible Cation Transport Regulator Homolog 1 (Chac1, also known as Botch) ^57, 58^ (**Figure 4d**). Chac1 is a γ-glutamyl cyclotransferase that negatively regulates Notch signaling in the mouse developing cortex ^59, 60^. Specifically, Chac1 is localized to the trans-golgi, where it prevents Notch signaling by interfering with the initial S1 furin-like cleavage step, keeping Notch in an immature, unprocessed form ^61, 62^ (**Figure 4e**). In line with the upregulation of Chac1 in Elp3cKO RGCs, we observed concomitant reduced levels of transmembrane (M) Notch1 (**Figures S5a-S5c**) in membrane fraction and cleaved Notch1 intracellular domain (NICD) (**Figure 4f**) in total extracts from P0.5 Elp3cKO lateral walls. Furthermore, the expression level of two canonical Notch signaling targets, *Hes5* and *Id3*, was reduced (**Figure 4g**), providing additional evidence that loss of Elongator activity impairs Notch signaling in ependymal cells of the lateral wall of Elp3cKO. A comparable induction of Chac1 was observed later, in P16 Elp3cKO lateral walls (**Figure 4h**), alongside with reduced NICD levels (**Figure 4i**) and Notch target gene expression (**Figure 4j**). Primary cilia length was already reduced in RGCs lining the third ventricle of E16.5 in Elp3cKO mice compared to WT littermates (**Figures S5d-S5g**). To determine whether restoring Notch signaling *in vivo* rescues ciliogenesis in differentiating RGCs, we depleted Elp3 in the RGCs by expressing Cre recombinase coupled to GFP via *in utero* electroporation of E16.5 Elp3^lox/lox^ mice (**Figure 4k**). Loss of Elp3 in RGCs resulted in significantly reduced cilia length at P0.5, a phenotype which was rescued by the concommitant expression of NICD (**Figures 4l-4m**). These data were further validated in human hTERT-RPE1 cells, a classical cell model to study primary ciliogenesis. These cells express GFP-tagged Arl13B to visualize primary cilia (**Figure S5h**). We confirmed a reduction in cilia length after Elp3 downregulation by esiRNA compared to esiRNA control (siSCR) (**Figures S5h-S5l**). This phenotype was associated with the upregulation of both *ATF4* and *Chac1* mRNA levels (**Figures S5j-S5k**). The supplementation of ISRIB (a blocker of the integrated stress response which also prevent ER stress transduction) or the expression of ATF4 siRNA (which prevents the transduction of ER stress downstream the activation of the Protein kinase RNA-like ER kinase (PERK)) rescued cilia length in siELP3 hTERT cells (**Figure S5l**). Notably, NICD overexpression in ELP3-downregulated hTERT cells also rescued cilia length (**Figure S5l**). As expected, Notch signaling-responsive genes *Hey1* and *Hes1* were downregulated in siELP3 hTERT cells, and NICD overexpression restored their expression levels (**Figures S5m-S5n**).

**Figure 4.**
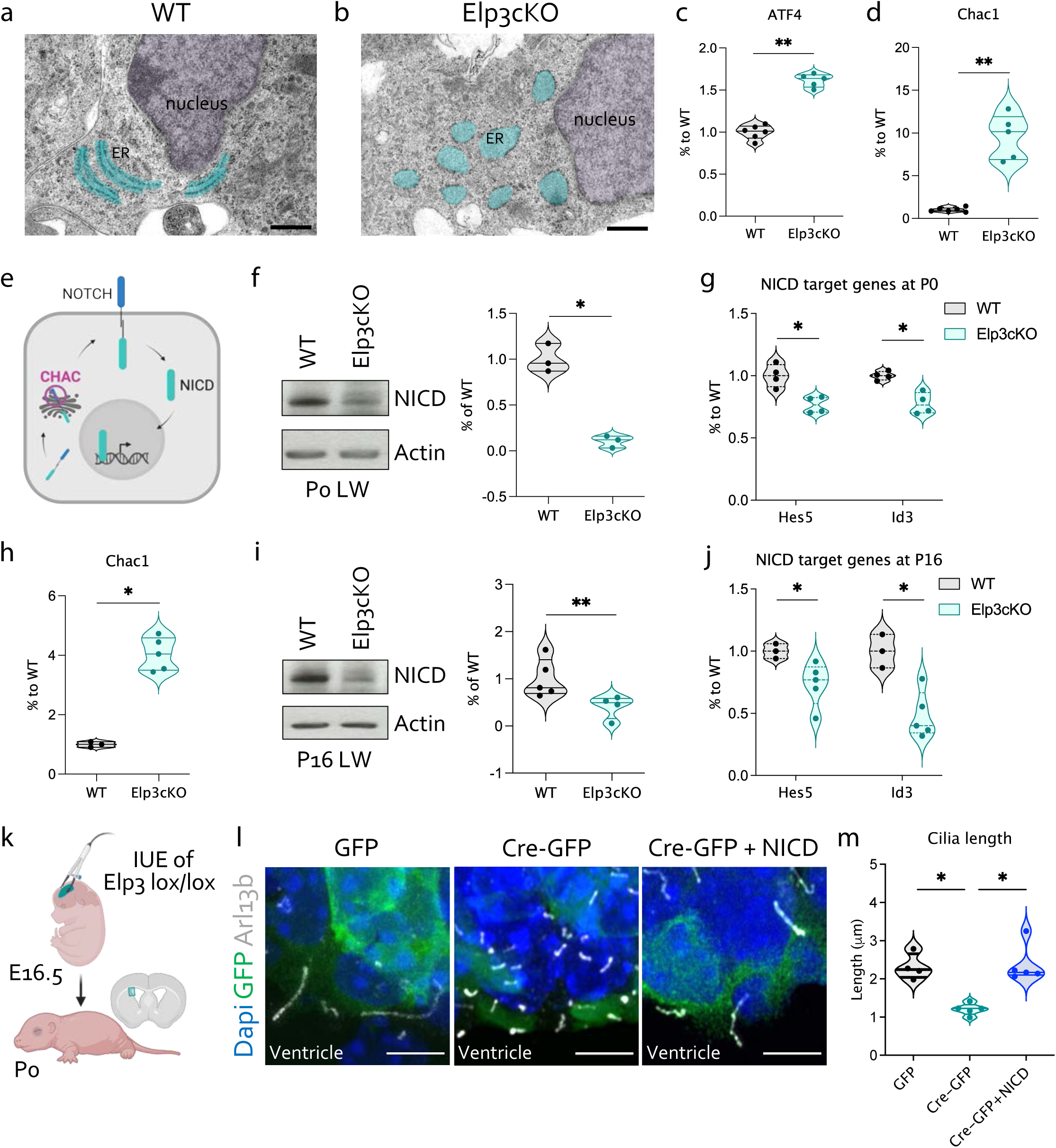
Elp3 depletion impairs primary ciliogenesis by interfering with Notch signaling. **a,b,** Transmission EM micrographs of WT (**a**) and Elp3cKO (**b**) RGC lining the lateral ventricle wall of P0.5 WT or Elp3cKO mice. The endoplasmic reticulum is blue highlighted and the nucleus is purple highlighed. Scale bars, 500nm. **c**, Violin plot of Atf4 mRNA expression with median ± quartiles from six WT and five Elp3cKO lateral walls. p-values were determined by Mann-Whitney test p=0.0043. **d,** Violin plot of Chac1 mRNA expression at P0.5 with median ± quartiles from six WT and five Elp3cKO lateral walls. p-values were determined by Mann-Whitney test p=0.0043. **e,** Scheme of a cell where Chac1 prevent the maturation of Notch1, thereby its membrane insertion and NICD release upon activation. **f,** Western blot of NICD from lateral wall extracts of P0.5 WT and Elp3cKO mice. Violin plot shows NICD at P0.5 with median ± quartiles from three WT and Elp3cKO lateral walls. p-values were determined by Mann-Whitney test p=0.05. **g,** Violin plots of hes5 and id3 mRNA expression with median ± quartiles from six WT and five P0.5 Elp3cKO lateral walls. p-values were determined by Mann-Whitney test p=0.0286 for both genes. **h,** Violin plot of Chac1 mRNA expression with median ± quartiles from three P16 WT and five Elp3cKO lateral walls. p-values were determined by Mann-Whitney test p=0.0357. **i,** Western blot of NICD on P16 lateral wall extracts of WT and Elp3cKO. Violin plot of NICD expression at P16 with median ± quartiles from five WT and four Elp3cKO lateral walls. p-values were determined by Mann-Whitney test p=0.0079. **j,** Violin plots of Hes5 and Id3 mRNA expression with median ± quartiles from three P16 WT and five Elp3cKO lateral walls. p-values were determined by Mann-Whitney test p=0.0357 for both genes. **k**, Scheme of the *in utero* electroporation strategy at E16.5 of Elp3^lox/lox^ mice to analyze cilia length at P0. **l,** Immunolabeling showing RGC cilia of electroporated cells expressing Arl13b (light grey), GFP (green) and Dapi (blue) upon expression of GFP only (control condition), cre-GFP (Elp3 depleted condition) and cre-GFP with NICD expressing plasmids (Elp3 is depleted together with NICD expression). Scale bars, 5µm. **m,** Violin plot of cilia length with median ± quartiles from three P0 WT and five Elp3cKO electroporated brains. p-values were determined by Kruskal-Wallis with Dunn’s multiple comparisons test p<0.05.

Altogether, these results indicate that ELP3 depletion leads to ER stress and upregulation of ATF4 and its direct target Chac1. This in turn impairs Notch1 signaling by preventing the membrane insertion of Notch1, which disrupts primary ciliogenesis in the differentiating RGCs that line the brain lateral wall.

### Loss of Elp3 leads to precocious differentiation of RGCs into ependymal cells

The blockade of Notch signaling arrests the proliferation of RGCs and guides their development into multiciliated ependymal cells across various organs and species ^17, 19, 63^. Here, we further investigated how RGCs differentiate into ependymal cells upon loss of Elp3 expression. We used dual pulse chase analysis upon successive intraperitoneal injections of pregnant dams with BrdU (bromodeoxyuridine) and EdU (ethynyldeoxyuridine) at embryonic day 16.5, with a 1.5-hour interval between the two pulses. Embryos were collected and fixed 30 minutes after the EdU pulse (**Figure 5a**).

**Figure 5.**
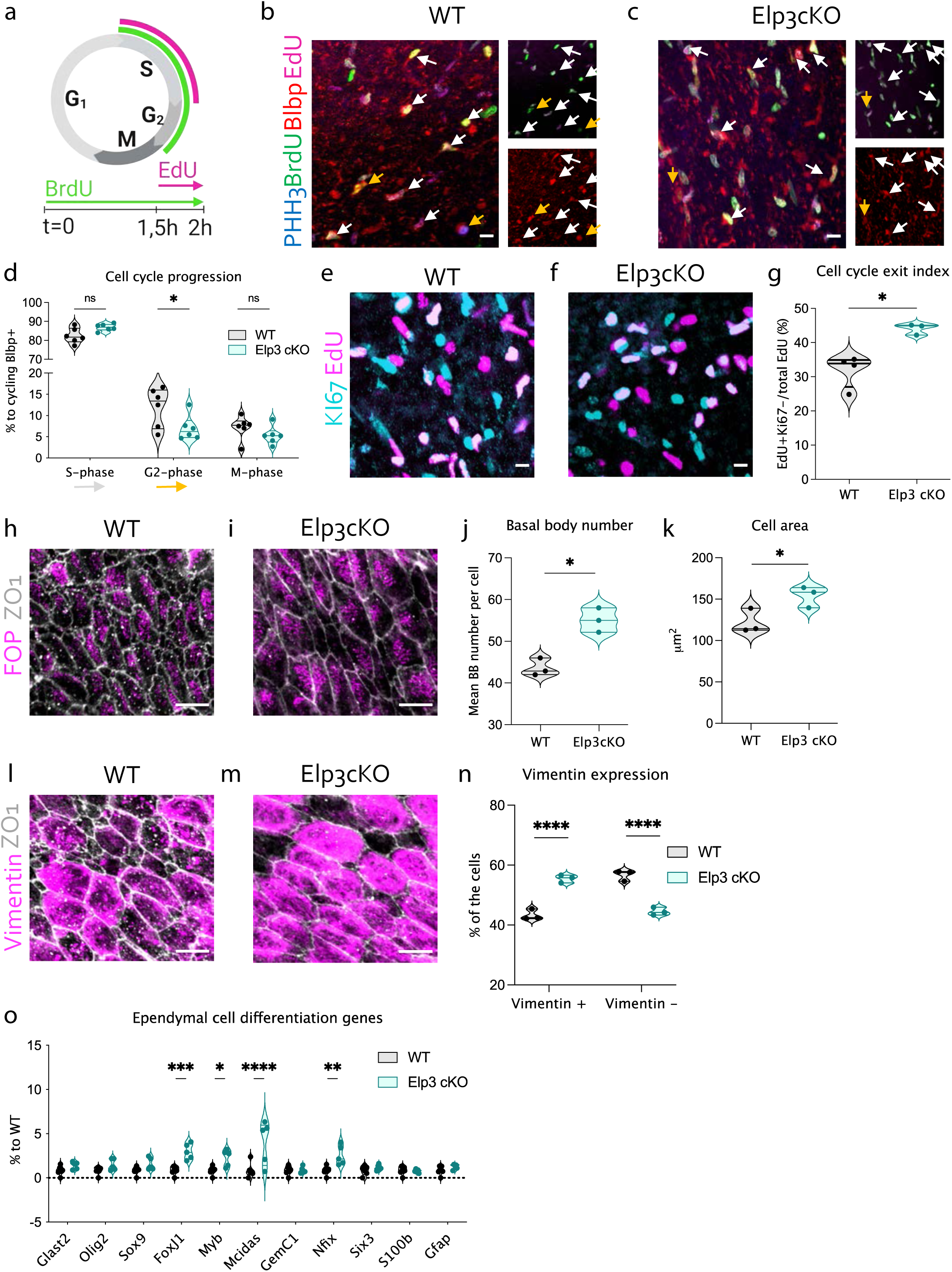
Loss of Elp3 leads to precocious differentiation of RGCs into ependymal cells. **a,** schematic representation of the dual-pulse labeling strategy with BrdU and EdU intraperitoneal injection of pregnant dams at embryonic day (E)16.5, with 1.5-hour interval between the injections. The embryos were collected and fixed 30 minutes after the EdU pulse. **b,c,** Immunolabeling of RGC with blbp (red), pHH3 (blue), BrdU (green) and EdU (magenta) on WT (**b**) and Elp3cKO (**c**) E16.5 lateral walls. Scale bars, 10µm. **d,** Violin plot of blbp^+^ RGCs in S-, G2- or M-phase of the cell cycle with median ± quartiles from six E16.5 WT and Elp3cKO lateral walls. p-values were determined by Two-way anova with Sidak’s multiple comparison test p<0.05. **e, f,** Immunolabeling of E16.5 WT (**e**) and Elp3cKO (**f**) embryos fixed 24 hours after the EdU pulse, labeled with KI67 (cyan) and EdU (magenta) to assess cell cycle exit. Scale bars, 10µm. **g,** Violin plot of the cell cycle exit index with median ± quartiles from four E17.5 WT and three Elp3cKO lateral walls. p-values were determined by Mann-Whitney test p=0.0285. **h,i,** Immunolabeling of cilia patches labelled with FOP (magenta) and ZO1 (light grey) of P7 WT (**h**) and Elp3cKO (**i**) lateral walls. Scale bars, 10µm. **j,** Violin plot of basal body number per cell with median ± quartiles of three P7 WT and three Elp3cKO lateral walls. p-values were determined by Mann-Whitney test p=0.05. **k,** Violin plot of cell area with median ± quartiles of three P7 WT and three Elp3cKO lateral walls. p-values were determined by Mann-Whitney test p=0.05. **l,m,** immunolabeling of ependymal cells labelled with vimentin (magenta) and ZO1 (light grey) of P7 WT (**l**) and Elp3cKO (**m**) lateral walls. Scale bars, 10µm. **n,** Violin plot of percentage of cells that are either vimentin positive or negative within a 100×100µm square with median ± quartiles of three P7 WT and Elp3cKO lateral walls. p-values were determined by Two-way anova with Sidak’s multiple comparison test ****p<0.0001. **o,** Violin plot of the percentage of mRNA expression detected in P7 Elp3cKO and WT lateral walls with median ± quartiles of six WT and five Elp3cKO biological replicates. p-values were determined by Two-way anova with Sidak’s multiple comparison test with *p<0.05, **p<0.01, ***p<0.001,****p<0.0001. BrdU, bromodeoxyuridine; EdU, ethynyldeoxyuridine

We analyzed the RGCs (Blbp^+^) labeled with BrdU and/or EdU along with a mitotic marker (pHH3) to assess the percentage of cycling RGCs that are in G2, S, or M phases (**Figures 5b-5c**). Although the percentage of cycling RGCs was not affected in Elp3cKO embryos (**Figures S6a-b)**, there was a significant reduction in the number of cycling RGCs in G2 phase compared with their WT littermates (**Figure 5d**). Moreover, we observed a complementary increase of the cell cycle exit index of RGC in Elp3cKO embryos (**Figures 5e-5g**).

The transformation of RGCs into ependymal precursors involves the amplification of centrioles, followed by their migration to the apical surface, attachment to the plasma membrane, and transformation into basal bodies that generate motile cilia ^1^. By P7, the differentiating Elp3cKO ependymal cells showed an increase in basal body numbers at the apical surface, along with a larger apical cell surface, indicating precocious ependymal cell differentiation as compared to their WT littermates (**Figures 5h-5k**). This came together with a higher percentage of differentiated vimentin-positive ependymal cells across the lateral wall of Elp3cKO (**Figures 5l-5n**). Furthermore, genes that controls ependymal cell differentiation, including *FoxJ1*, *c-Myb*, *Mcidas*, and *Nfix*, were expressed at higher levels in Elp3cKO P7 lateral wall as compared to WT littermates (**Figure 5o**). Overall, these findings suggest that the disruption of Notch signaling in Elp3cKO RGCs leads to increased cell cycle exit and early ependymal cell differentiation.

### Postnatal depletion of Elp3 does not affect translational and rotational polarities

During the course of lateral wall maturation, establishment of RGC polarity precedes and is required to establish ependymal cell polarity ^9, 23^. To assess wether defects observed in mature Elp3cKO lateral walls stem from early defect, we used a tamoxifen-inducible conditional knock-out mouse model to deplete Elp3 postnatally in postmitotic RGCs in the process of ependymal cell differentiation. This was achieved by breeding Elp3^lox/lox^ mice with Glast-CreERT2 transgenic mice and R26tdTomato reporter mice, hereafter referred to as Elp3icKO. To induce Elp3 deletion, we injected tamoxifen into dams from P1 to P4, which was passed to the pups via breast feeding. Reduced Elp3 mRNA expression was confirmed in FACS-purified tdTomato^+^ cells from P7 Elp3icKO pups as compared to those isolated from their WT littermates (**Figures 6a-6b**). The expression level of key ependymal cell differentiation genes was unchanged between Elp3icKO and WT animals (**Figure 6c**). Next, we evaluated rotational (**Figures 6d-6j**) and translational polarities (**Figures 6k-6r**) at P16, as described above, and did not observed any significant differences between Elp3icKO and WT, demonstrating the conservation of organized polarity. These results indicate that the early postnatal depletion of Elp3 does not impact the proper maturation of ependymal cells.

**Figure 6.**
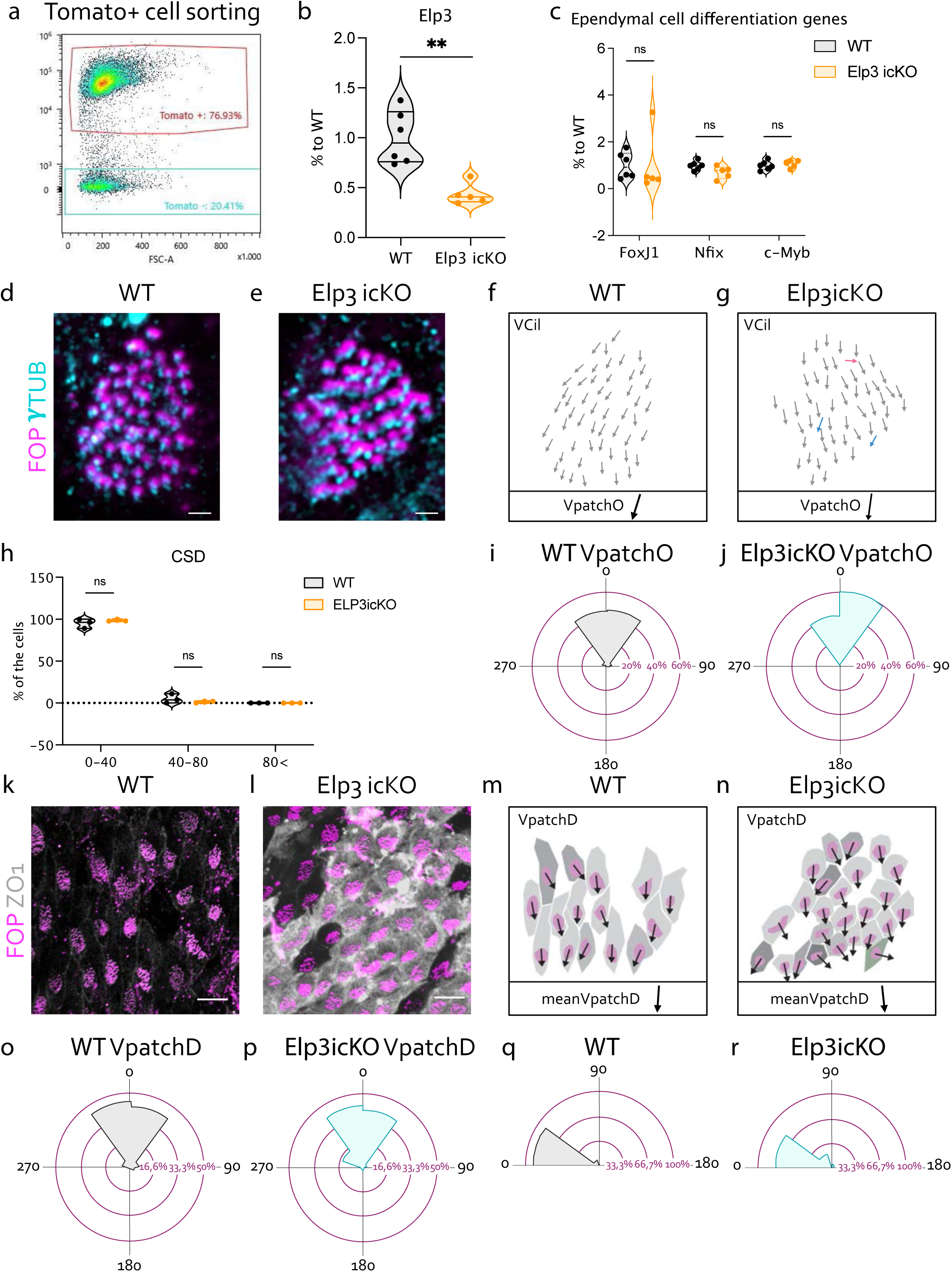
Postnatal depletion of Elp3 does not affect translational and rotational polarities. **a,** FACS-gating strategy to isolate tdTomato^+^ ependymal cells from P7 lateral walls. **b,** Violin plot of Elp3 mRNA expression in six P7 WT and five Elp3icKO lateral walls. p-values were determined by Mann Whitney test p=0.0043. **c,** Violin plot of the percentage of mRNA expression detected in Elp3icKO tdTomato^+^ ependymal cells, as compared to their WT controls with median ± quartiles of six P7 WT and five Elp3cKO lateral walls. Two-way ANOVA with Bonferroni’s multiple comparisons test revealed no statistical differences between groups. **d,e,** Immunolabeling of motile cilia patches labelled with FOP (magenta) and γ-tubulin (cyan) on WT (**d**) and Elp3icKO (**e**) on the dorsoventral region of the lateral wall. Scale bars, 1µm. **f,g,** Ciliary vector was drawn according the immunostainings, representing the polarity of each cilium within the patch for WT (**f**) and Elp3 icKO (**g**). Colors representing the different ciliary vector directions. VpatchO represents the mean of the individual ciliary vectors. **h** Percentage of ependymal cells according to the ciliary vector standard deviation (CSD) in WT and Elp3icKO P16 lateral wall. Violin plots show median ± quartiles from three biological replicates. Two-way ANOVA with Bonferroni’s multiple comparisons test revealed no statistical differences between groups. **i,j,** Circular dispersion plots of VpatchO around the mean in WT (216 cells) (**i**) and Elp3 cKO (114 cells) (**j**) from three biological replicates. p-values were determined by Watson’s U^2^ test= 0,133; ns. **k,l,** Confocal images of cilia patches labelled with FOP (magenta) and ZO1 and tomato (light grey) and on WT (**k**) and Elp3icKO (**l**) lateral wall. Scale bars, 10µm. **m,n,** A vector indicating the displacement of the BB patch from the geometrical center of each cell’s apical surface to the center of the BB patch was drawn according the immunostainings for WT (**m**) and Elp3icKO (**n**). Pseudocolors representing the different BB patch displacement directions. VpatchD represents the mean of the individual displacement vectors. **o,p,** Representation of the circular dispersion of individual VpatchD around the mean in WT and Elp3cKO; brain lateral wall area from three biological replicates. p values were determined by Watson U^2^ test= 0,247; ns. **q,r,** Circular dispersion of the correlation between VpatchD (translational polarity) and VpatchO (rotational polarity) angles in WT and Elp3cKO; from three biological replicates. p values were determined by Watson U^2^ test= 0,446; p<0.001***.

## Discussion

Here we show that the loss of Elongator activity in RGCs via Elp3 depletion disrupts the generation and maturation of the ependymal cells that line the brain’s lateral wall. This phenotype results from the induction of ER stress together with the upregulation of ATF4 and its direct target Chac1 that reduces the intracellular maturation of Notch1 receptors, further preventing its proper membrane insertion in RGCs. As a result, there is a shortening of the primary cilium, along with an early exit from the cell cycle and an accelerated maturation of RGCs into ependymal cells. This shift in maturation timing is accompanied by significant changes in PCP-related processes, which are crucial for the functional organization of motile cilia and CSF circulation.

Elp3 is the catalytic subunit of the Elongator macromolecular complex, which consists of six subunits (Elp1–Elp6) ^33^. This highly conserved complex is essential for neurodevelopment, and mutations in the genes encoding its subunits have been linked to intellectual disability and various neurological disorders ^64^. At the molecular level, evidences from yeast and mouse models have demonstrated that the Elongator complex plays a critical role in the modification of wobble uridine of selected tRNAs, thereby directly influencing RNA translation efficiency ^34, 38–43^. A recent study using a mouse model with conditional depletion of Elp1 under the Tuba1a promoter demonstrated kyphosis and microcephaly accompanied by enlarged lateral ventricles^65^. The authors reported a reduction in the number of RGC primary cilia protruding into the ventricle at E14.5, as well as a decreased number of ependymal cells with cilia tufts in the adult ventricle. Here, we demonstrated that the conditional deletion of Elp3 in forebrain progenitors affects the cilia length of RGCs at E16.5, and is further followed by a severe disorganization of cilia tufts of ependymal cell in the lateral wall postnatally. Interestingly, in addition to the cilium phenotype, we observed a marked reduction in the number of microvilli at the surface of ependymal cells across the lateral wall surface in Elp3cKO P0.5 pups. Microvilli are extracellular protrusions enriched with bundled actin filaments^66^, and we have previously demonstrated that the loss of Elp3 activity leads to actin filament severing through increased cofilin activity^35^. It would be worthwhile to investigate whether the decreased density of microvilli at the surface of ependymal cells observed in Elp3cKO mice results from such defect. Here, we demonstrate that Elp3 is essential at the single-cell level for the intrinsic organization of cilia patches, as its loss results in disorganized cilia tufts in ependymal cells on the lateral wall surface. Although the cilia motility of ependymal cells itself is not impaired in Elp3cKO, our data suggest that defects in rotational and translational polarities lead to beating defects and disrupted directed ventricular fluid flow, consequently contributing to the development of postnatal hydrocephalus. The asymmetric distribution of planar cell polarity (PCP) proteins is crucial for the unified alignment of BBs within individual ependymal cells and across the entire ependyma. Mutations in core PCP genes have been shown to disrupt rotational polarity (e.g., *Celsr2/3*, *Vangl2*, and *Dvl2*) and impair translational polarity (e.g., *Celsr1*, *Fzd3*, and *Vangl2*) in ependymal cells ^23, 25–27, 67^. Interestingly, we found that Elp3 deletion did not alter the expression levels of core PCP genes or the typical localization of Vangl2 on the opposite side of the ciliary patch in maturating ependymal cells. However, we observed a disrupted cellular distribution of Vangl2 at the tissue level. This indicates that the uncoordinated distribution of Vangl2 across the ependymal tissue is likely a consequence of upstream events triggered by the loss of Elp3 in RGCs. Additionally, our data demonstrate that, although the BBs of RGCs undergo lateral displacement in Elp3cKO P0.5 lateral walls, the coordinated displacement of BBs between neighboring RGCs is disrupted. This suggests an early loss of intercellular coordination of translational polarity in RGCs. These findings align with the current understanding of ependymal planar cell polarity establishment as a multistep process, initially guided by the translational polarity of primary cilia in RGCs and subsequently refined by motile cilia in ependymal cells, as originally proposed ^9^.

Further ultrastructural analysis of the primary cilium and its anchorage to the apical surface from the basal body confirmed the shortening of the cilium, accompanied by a slight accumulation of electron-dense material in the central core of the cilium in Elp3cKO ependymal cells. This latter phenotype has not been reported previously and provides novel insights into the structural alterations associated with Elp3 deficiency. Notably, this accumulation is unlikely to result from the accumulation of motile vesicles along axonemal MTs due to impaired intraflagellar transport, which would typically appear at the periphery of the primary cilium. Previous studies have demonstrated that both actin and MTs are crucial for primary cilium maturation, centriolar migration, docking, and proper spacing ^68–70^. BBs are transported to the apical surface of ependymal cells through an actomyosin-driven mechanism ^71^. Although Elp3 has been reported to modulate actin and MT dynamics in migrating and differentiating cortical neurons ^35, 38, 72^, our findings did not indicate disrupted BB migration in Elp3cKO animals. Instead, we observed an increased number of BBs within ependymal cells, suggesting precocious differentiation. The loss of Elongator activity has been previously associated with reduced MT acetylation in migrating projection neurons and impaired bidirectional axonal transport of vesicles, which are critical for MT acetylation in axons ^72^. However, we did not detect changes in MT acetylation levels in the LWs of P16 Elp3cKO mice.

The molecular coupling between ciliogenesis and PCP signaling remains debated. Indeed, some studies suggest that PCP may operate independently of ciliogenesis. For instance, in mouse models with conditional ablation of Kif3a in motile cilia, rotational polarity is disrupted while translational polarity remains unaffected ^9^. Furthermore, the p73 isoform TAp73 is both necessary and sufficient for the membrane localization of PCP-core proteins in a non-ciliated cellular model, indicating that the regulation of PCP by p73 can occur independently of its function in multiciliogenesis ^73^. Conversely, the loss of expression of certain PCP-core components impairs ciliogenesis in ependymal cells ^23^. Furthermore, analyses of Vangl and Celsr mutants indicate that, while PCP signaling regulates cilia positioning, it does not directly influence the ciliogenesis process itself. In our study, we observed that the postnatal depletion of Elp3 does not disrupt the establishment of translational and rotational polarity in ependymal cells. Our finding suggests that the anatomical and functional disorganization of motile cilia in Elp3cKO ependymal cells arises from primary ciliogenesis defects that take place earlier in RGCs.

Notch signaling is key for the regulation of both ciliogenesis and cell cycle progression. For example, the Kupffer’s vesicle, a transient structure of the developing zebrafish, harbours motile cilia whose development relies on Notch signaling. Specifically, upregulation of Notch increases ciliary length, while its downregulation results in shorter cilia. These variations in ciliary length significantly influence the flow within the Kupffer’s vesicle, thereby affecting the development of left-right asymmetry^74^. Dorso-ventral patterning of the neural tube in both mouse and chicken models also requires ciliary extension dependent on Notch signaling. In this context, NICD modulates the cellular response to Sonic Hedgehog by enhancing both ciliary length and the localization of Smoothened within the cilia ^75^. This aligns well with our data, which demonstrate that disrupting NICD signaling affects the length of primary cilia in RGCs, a process that is disrupted upon loss of Elp3 expresion. This phenotype is driven by the upregulation of ATF4, which in turn promotes the expression of Chac1. Chac1 inhibits the maturation of Notch1 required for its membrane insertion, thereby disrupting NICD signaling. The ultrastructural analysis of perinatal RGCs lacking Elongator activity reveals a significant enlargement of ER tubules, which is a hallmark of ER stress. Although ER stress has been linked to the activation of the UPR-associated PERK/ATF4 pathway in cortical progenitors ^76^, we cannot rule out the possibility that a portion of ATF4 may also be stabilized through mTORC1 activation, as previously observed in intestinal progenitors ^77^.

The length of the primary cilium has also been linked to cell cycle progression. Depletion of Nde1 delays G0/S transition, which correlates with an increase in primary cilium length ^78^. Similarly, in Elp3cKO mice, RGCs exhibit shorter primary cilia, which corresponds to their early cell cycle exit and premature differentiation into ependymal cells. Recent studies have linked genes involved in cell cycle progression to the multiciliation cycle, which drives terminal differentiation following progenitor cell division ^79, 80^. Four distinct phases within the multiciliation cycle, analogous to those of the cell cycle, have been identified: amplification (S-like phase), growth and disengagement (G2/M-like phase), and ciliogenesis (G1/G0-like phase). E2F7 has been shown to be critical for the proper termination of the S-like phase and for facilitating the transition to the G2/M-like phase, thus influencing centriole maturation and subsequent ciliogenesis. In multiciliated cells lacking E2F7, centrioles remain undocked, and ciliogenesis is diminished ^79^. Our RT-qPCR analysis of the lateral wall from Elp3cKO mice did not show a reduction in E2F7 gene expression at P0 or P7 (data not shown), further supporting the hypothesis that the loss of Elongator does not lead to premature differentiation of ependymal cells by disrupting the multiciliation cycle ^79^.

All data combined within this study adds to the growing understanding of the critical function for the Elongator complex in the CNS by revealing that the loss of Elp3 in the developing cortex caused the hyperactivation of ER stress in RGCs resulting in defective maturation of Notch receptors, ultimately disrupting the timing of ependymal cell maturation.

## Material and methods

### Mouse genetics

All mice were handled according to the ethical guidelines of the Belgian Ministry of Agriculture in agreement with European Community Laboratory Animal Care and Use Regulations (86/609/CEE, Journal Officiel des Communautés Européennes, L358, 18 December 1986) and with our experimental protocol n°21-2443. Both males and females were used for all analyses. FoxG1-Cre (kindly provided by J.-M. Hébert) and Elp3^lox/lox^ mice were described previously ^34, 81^. Glast-creERT2 ^82^ crossed with R26CAGTdTomato (The Jackson Laboratory, stock number 007914) and Elp3^lox/lox^ mice were used as a transgenic model to characterize the loss of Elp3 postnatally.

### Western blot

Dissected P0 and P16 lateral walls were lysed in lysis buffer (50 mM Tris-HCl pH 7.4, 450 mM NaCl, 1% Triton X100, 10 mM NaF, 1 mM Na3VO4 and Complete Proteases Inhibitors Cocktail, #04693116001 Roche). Proteins were extracted by centrifugation (10,000 g) for 10 minutes at 4°C, and quantified using BCA method (#23225 Pierce). Protein separated on 8% SDS-PAGE gels. For tubulin and acetyl-tubulin quantifications, only two microgram of protein were loaded. Nitrocellulose membranes (# 88018, Thermo Scientific) were incubated O/N at 4°C with primary antibodies including, Elp3 (1:1000, #5728, CST), α-tubulin (1:10000, #ab19251, Abcam), acetylated α-tubulin (1:10000, #T7451, Sigma), cleaved notch1 (1:500, #4147, CST), Notch1 (1:1000, #sc-9170, SantaCruz), actin-HRP (1:5000, #3865, Sigma). After washing, secondary antibodies conjugated with horseradish peroxidase (1:1000 #G-21234, ThermoFisher, for primary antibodies raised from rabbit; 1:1000, #G-21040, ThermoFisher, for primary antibodies raised from mouse) were applied for one hour at RT. The signals were visualized with ECL chemiluminescent reagent (#34580, Thermo Fisher Scientific) using either Hyperfilm ECL (#GE28-9068-35, Merck) or by ImageQuant800 Amersham (cytiva).

### Cytosolic and membrane fractionation

Dissected LW were lysed in lysis buffer (250mM sucrose, 50mM Tris-HCl, 5mM MgCl2, 1mM DTT and Complete Proteases Inhibitors Cocktail (#04693116001, Roche) by sonication on ice (3 times 10 s bursts with intensity ∼40% and 30 s breaks) and centrifuged for 15minutes at 800g to remove nuclei. The supernatant 1 was centrifuged again at 1,000g for 15 min. The obtained supernatant 2 was subjected to an ultracentrifugation step for 1 h at 100000g to obtain the cytosolic and membrane fractions. The supernatant contained the cytosolic fraction and the pellet was incubated in lysis buffer 2 (20 mM Tris-HCl, 0.4 M NaCl, 15% glycerol, 1.5% Triton X-100, 1mM DTT and Complete Proteases Inhibitors Cocktail) for 1 hour shaking at 1400 rpm and 4 °C and centrifuged for 30 minutes at 9000g of which the supernatant contained the membrane proteins.

### Cell line

hTERT-RPE1 stably express Arl13b-GFP protein (gift from Dr.F. Saudou, Institut des Neurosciences, Grenoble, France)^83^. Cells were transfected with 15pmol mission esiRNA against Elp3 or Firefly Luciferase as control (#EMU087711 and #EHUFLUC, respectively, Sigma) using lipofectamine 2000 transfection reagent (#11668019 ThermoFisher). After 72h of transfection, cells were serum starved for 48h in opti-MEM® medium (#31985070, Gibco) and fixed in 4% paraformaldehyde (PFA) for immunohistochemistry analysis (as described below).

### Immunohistochemistry

Dissected LWs were fixed 12 minutes in 4% PFA containing 0.1% triton X-100, washed three times in PBS and incubated 4hours in PBS 0.1% Triton X-100, 10% donkey serum at 4°C. LWs were successively incubated 48hours at 4°C with primary antibodies including, γ-tubulin (1:300; #T6557, Sigma), anti-ZO1 (1:300, #ab19251, Abcam or #14-9776-82, Invitrogen), anti-FGFR1OP (1:300, #H00011116-MO1, Abnova), Vangl2 (1:250, gift from Mireille Montcouquiol), Arl13b, Notch1 (1:200, #sc-9170, SantaCruz), acetylated α-tubulin (1:1000, #T7451, Sigma), GFP (1:500, #ab6673, Abcam), Blbp (1:300, #ABN14, Millipore), KI67 (1:500, #AB9260, Millipore), Vimentin (1:500, #ab24525, Abcam). LWs were incubated 24hours at 4°C with secondary antibodies (1:500, ThermoFisher). Nuclei were counterstained with DAPI (1:5000, #, Sigma). Five non-overlapping fields (50×50µm for P0 and 100×100µm for P16) were acquired for each sample on a confocal microscope NikonA1R using a 60X oil objective or ZEISS LSM980 using a 63X oil objective. For immunolabeling on P7 coronal slices, slices were treated with 10mM citrate buffer (pH6) supplemented with 0.1%tween20 for 10 minutes at 95°C, blocked in 10% donkey serum in PBS-tween 0.1% for 4hours at 4°C and primary antibodies were incubated overnight at 4°C, including Elp3 (1/200; #17016-1-AP, proteintech) Arl13b (1/250, #75-287, Antibodies Incorporate) and FoxJ1 (1/200, #149965-82, Invitrogen) Slices were incubated 2hours at RT with secondary antibodies (1:500, ThermoFisher). Images were acquired on a confocal microscope (ZEISS LSM980) using a 63X oil objective.

### Dual pulse labeling

Pregnant dams were injected at E16.5 gestational day following a dual pulse labeling method with bromodeoxyuridine (BrdU, #B23151 ThermoFisher) and ethynyldeoxyuridine (EdU, (#C10337, Invitrogen) with a 1.5-hour interval between the two pulses, and fixed the embryos in 4% PFA at RT 30 minutes after the EdU pulse. Immunolabeling was performed as above with some minor adaptations. Prior to the blocking of 4hours in PBS 0.1% Triton X-100, 10% donkey serum at 4°C, the E16.5 LW were subjected to an antigen retrieval step of two times 30 minutes 2N HCl at 37°C and consequent incubation of 0.1 M sodium borate pH8.5 for 10minutes at RT. LWs were successively incubated 48hours at 4°C with primary antibodies including, BrdU (1:300, #MCA2060, AbD Serotec), pHH3 (1:500, #MA5-15220, ThermoFisher), Blbp (1:300, #ABN14, Millipore). LWs were incubated 24hours at 4°C with secondary antibodies (1:500, ThermoFisher). EdU revelation was performed with the Click-iT™ EdU cell proliferation kit for imaging (#C10337, Invitrogen) according the manufacturer’s instructions after the secondary antibody incubation.

### Fluid Flow Analysis

Freshly dissected lateral walls from WT or Elp3cKO animals were immobilized in Petri dishes containing Leibovitz media at 37 °C. Hand-pulled glass capillaries were used to release a solution containing 5% GFP-fluorescent microbeads (2-µm diameter; #09847; Biovalley) and 5% glycerol in water at the anterior dorsal part of the lateral wall. Bead movements were recorded using an iPhone 6 mounted on a Leica M80 microscope. Several rounds of beads release were performed for each ventricle during a maximum of 15 minutes. During the course of the experiment, beads progressively dropped along the flow pathway. At the end of recordings, non-deposited beads remaining in the medium were removed, and deposits of beads on the roof of lateral ventricle were photographed.

### RNA labeling

Co-detection Basescope (#323910 and #323180, Biotechne) according manufacturers instructions, with minor adaptations. Primary antibody (ZO1 1/200 rabbit) was incubated overnight at 4°C on the slides before the basescope protocol with four minutes of co-detection target retrieval, twelve minutes of protease Plus incubation and secondary antibody (1/300) with Dapi (1/1000) were put overnight at 4°C. After washing the slides were mounted and images on a ZeissLSM980. Elp3 probe (BaseScopeTM Probe - BA-Mm-Elp3-E2-1zz-st-C1,Mus musculus elongator acetyltransferase complex subunit 3 (Elp3) transcript variant 1 mRNA) was designed by Biotechne.

### In utero electroporation

*In utero* electroporation was performed on E14.5 timed embryos as described previously ^84^. The pregnant mice were injected with buprenorphine (Temgesic, Schering-Plough, Brussels, Belgium), deeply anaesthetized with isoflurane in oxygen carrier and placed on heating pad. The uterine horns were exposed through a 1.5 cm incision in the ventral peritoneum. Plasmid used were: pCIG2-IRES-GFP (3µg/µL); pCIG2-Cre-IRES-GFP (3µg/µL), pCAG backbone (2µg/µl), pCAG-NICD (2µg/µL). The plasmid solutions were injected through the uterine wall into the lateral vesicle using a Femtojet microinjector (VWR). Five electrical pulses (45 V, 50 ms duration, 1 sec off) were delivered across the head of the embryo (targeting the dorsal-medial part of the cortex) using 5 mm platinum tweezers electrodes (CUY650P5, Sonidel, Ireland) and ECM-830 BTX square wave electroporator (VWR). The uterine horns were then replaced in the abdominal cavity and the abdomen wall and skin were sutured using surgical needle and thread. The pregnant mouse was placed under a warming red light until complete awakening.

### Planar cell polarity analysis and quantification

Analyses and quantifications were performed using software Biotool1 (github.com/pol51/biotool1). For analysis of γ-tubulin and ZO1 immunostainings, the contours of ependymal cells and BBs patches were manually traced. The software was used to calculate (*i*) the geometric center of these contours, (*ii*) to analyze the coordination of patch displacement, VpatchD defined from the center of the cell to the center of the patch, and the deviation of individual vectors relative to the mean vector of the field meanVpatchD. A similar strategy was used to evaluate the coordination of the primary cilium displacement (VD and meanVD). In that case, a vector was drawn between the center of the cell and the primary cilium BB. Cells where the BB is superposed to the center were taken into account.

For analysis of ciliary patches using FOP and γ-tubulin double immunostaining, dots corresponding to both signals were manually defined for each cilium. A unit vector connecting the two dots was defined by the software (Vcil). These vectors were used to calculate the CSD of each patch. The distribution of cells in the different bins of CSD was done in Excel. The coordination of patch orientation was analyzed from the same manual acquisition. Only cells with CSD below 40° were considered to define VpatchO (meanVcil). The deviation of individual VpatchOs relative to the meanVpatchO was calculated to evaluate the tissue coordination of the patch orientation.

To analyze the intracellular coordination of VpatchD and VpatchO, cells, patches, and Vcils were defined after manual tracing on ZO1, FOP, and γ-tubulin triple immunostaining. Angles between individual VpatchDs and VpatchOs were calculated.

The output of Biotool1 is a list of angles that were plotted in Oriana software (Kovach Computing Services) to obtain the graphical representation of their distribution in 30° bins and perform circular statistical analysis. In all circular representations, the percentage of cells is represented as the radius of wedge. For statistical analysis, the Watson U^2^ test for analyzing 2 samples of circular data was applied.

### Electron Microscopy Transmission Electron Microscopy

The brains were dissected and fixed at 4°C in 2.5% glutaraldehyde in 0.1 M Sörensen’s buffer (0.2 M NaH2PO4, 0.2 M Na2HPO4, pH 7.4) for 2 hours, then postfixed at 4 °C with 2% osmium tetroxide in Sörensen’s buffer for 1 hour, dehydrated (70%, 96% and 100% ethanol) at room temperature and embedded in Epon for 48 hours at 60 °C. Seventy nanometer sections were obtained (Reichert Ultracut E) and mounted on copper grids coated with collodion. The sections were contrasted with uranyl acetate and lead citrate for 15 minutes each. The sections were analysed under a JEM-1400 transmission electron microscope (Jeol) at 80 kV and pictures were obtained with an 11 MegaPixel bottom-mounted TEM camera system (Quemesa, Olympus).

### Scanning Electron Microscopy

Dissected LW were fixed at 4°C 2hours in 2.5% glutaraldehyde in 0.1 M Sörensen’s buffer, then post-fixed with 1% osmium tetraoxide 1hour at 4°C. The specimens were dehydrated, critical point dried in CO2, mounted on sputter grids coated with platinum, and analyzed on a FEI ESEM-FEG XL-30scanning electron microscope (Philips).

### Fluorescent Activated Cell Sorting (FACS)

LW were dissected and processed to obtain a single cell suspension by incubating them for 20 minutes at 37°C in 28 U/mL papain (#LK003176, Worthington) supplemented with 0,05% DNAse1 (#D5025, Sigma). Live cells were gated as dapi negative (405nm laser) using the scatter gate, doublets were excluded and tomato+ cells (561nm laser) were retrieved on a SonyMA900 sorter.

### RNA extractions and qRT-PCR

Total RNA was extracted from microdissected LW at P0, P7 and P16, HTERT ARl13b-GFP cells after 48h of serum starvation or from tomato+ FACS-sorted cells. RNA extraction was performed with TRIzol Reagent (#15596018, Invitrogen) according to the manufacturer’s instructions, then treated with DNAse I (#AM1907, Invitrogen). cDNA was synthesized using the RevertAid H Minus First Strand cDNA Synthesis kit (#K1632, Thermo Fisher). Quantitative real-time PCR was performed in duplicates using SYBR Green (#A6002, promega) on a LightCycler 480 System (Roche). A standard curve was used to calculate relative concentrations of gene expression per gene. An average of technical duplicates was made, and normalized to the average of the 3 HKGs (GAPDH, PPiA, and 36B4). Data were analyzed with LightCycler 480 software version 1.5.1.62 (Roche Life Sciences) and Microsoft Excel, and graphs were made in Prism 10 (GraphPad Software).

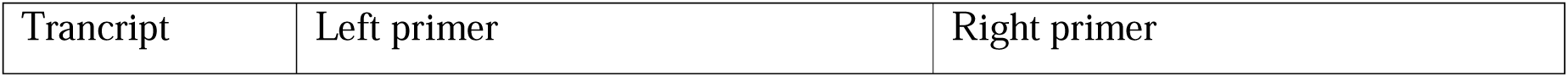
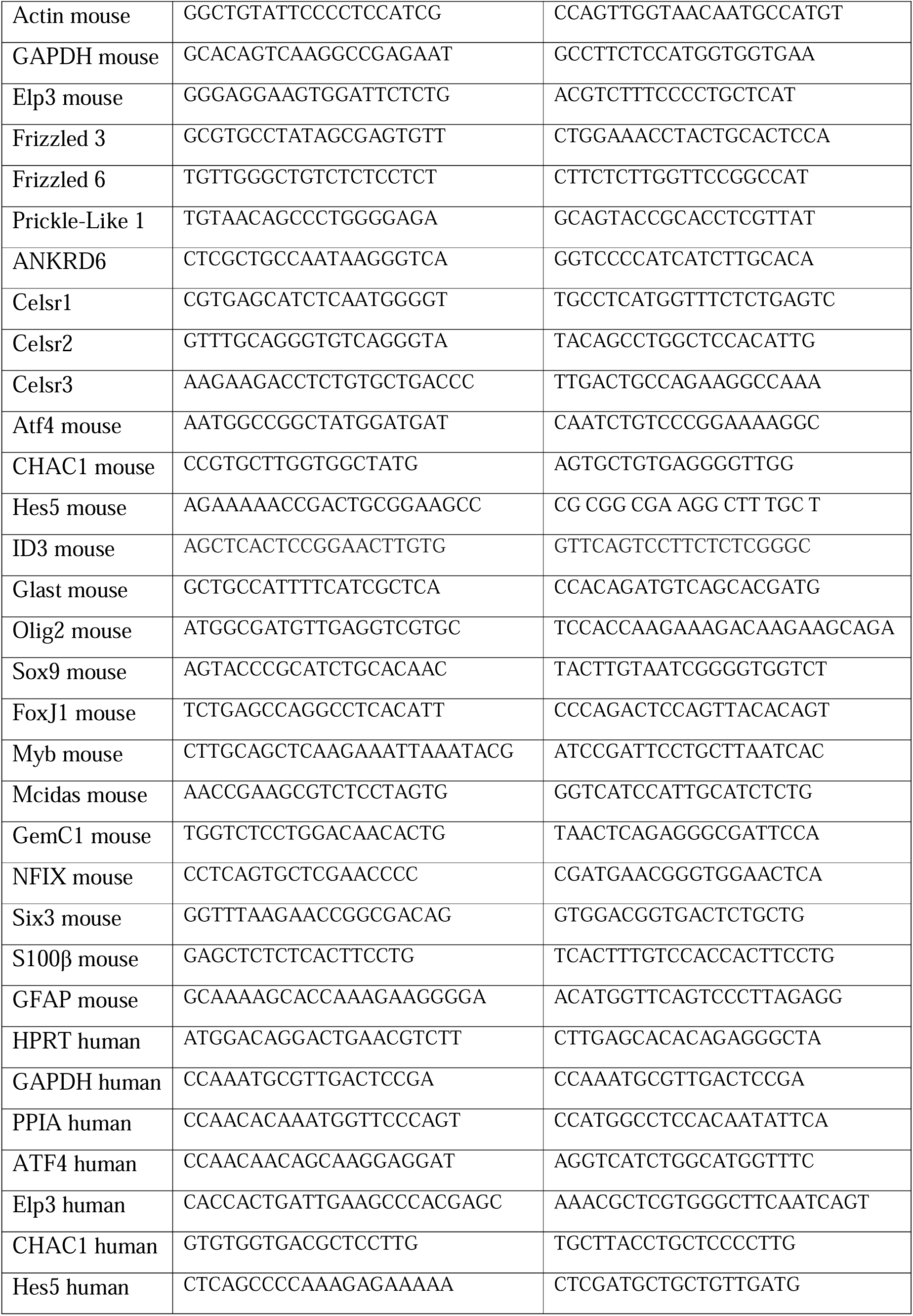
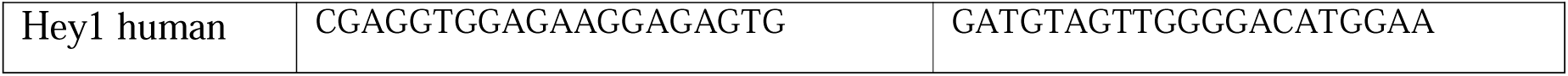

### Data collection and curation

Single cell RNA-seq (scRNA-seq) dataset was collected from the GEO database (<u>GSE123335</u>, specifically the P0 combined matrix and cluster annotations). The dataset was analyzed using the Seurat R package, version 4.3.0. Filters, adjusted to the dataset, were chosen to keep cells with less than 3000 unique features, 6500 RNA counts and[6% mitochondrial RNA. The data was normalized with the SCTransform function, regressing the percentage of mitochondrial genes. PCA was run and the 30 first principal components were used as dimensions to perform an UMAP non-linear dimensional reduction. Finally, the nearest neighbor graph was generated using 25 dimensions and the clustering was performed using algorithm 1 with a resolution set to 0.5. Clusters were identified and grouped together based on the expression of one or multiple marker genes previously identified ^44, 85–88^.

| Cluster | Marker genes |
| --- | --- |
| Apical progenitors | *Pax6, Hes5, Aldoc, Ddah1* |
| Intermediate progenitors | *Eomes* |
| Interneurons | *Dlx1, Dlx2, Gad1, Gad2* |
| Cajal retzius cells | *Lhx1, Reln, Trp73* |
| Projection neurons | *Neurod1, Sema3c, Neurod2, Neurod6* |
| Oligodendrocytes precursor cells | *Olig1, Olig2, Pdgfra, Sox10* |
| Astrocytes | *Aldh1l1, Aqp4, Sparcl1* |
| Endothelial cells | *Cldn5, Igfbp7* |
| Microglia | *Aif1, C1qb, C1qc* |
| Perycites | *Vtn, Pdgfrb, Kcnj8* |
| Ependymal cells | *Foxj1, Pifo, Dynlrb2* |

### Statistical Analyses

Statistical information, including n, mean and statistical significance values, are indicated in the figures or figure legends. Data are plotted using Graphpad Prism 10 and for circular statistics, WT and Elp3 cKO mutants were compared using the Watson U2 test (Oriana software). N (number of biological replicates) is represented in the figures or figure legends and statistical tests were performed on means ± s.d. of biological replicates. At least three biological replicates were used per experiment. For quantification of the ciliary length, microvilli surface of microdissected LWs, on coronal slices and of HTERT arl13b-GFP cells, ImageJ was used. Statistical significance was determined with Graphpad Prism 10 using the tests indicated in each figure. Data were considered statistically significant at *P[<[0.05, **P[<[0.01, ***P[<[0.001 and ****P[<[0.0001.

## Acknowledgments

The authors thank Dr. Hébert for providing Cre-mouse line; Dr. F Saudou for providing HTERT-RPE1-arl13b-GFP cells. Dr. Prevot, G. Duysens and N. Kruzy for technical assistance. We thank Drs B. Franco, L. Broix, M. Kuchma for their scientific input. We thank the technological GIGA platforms (Cell imaging, Flow cytometry, and Viral vectors) for providing tools and assistance to acquire experimental data. We thank Florian Loria for graphical assistance. Some schematic representation were designed with BioRender (https://www.biorender.com/). S.T. is supported by Win2Wal (ChipOmics; #2010126). M.C-B is supported by the Marguerite-Marie Delacroix fund; L.O., C.R., A.B., S.M.S are supported by the F.R.S-FNRS; C.C. is a postdoctoral researcher from the F.R.S-FNRS; L.N., A.C, and B.M are Research Director from F.R.S-FNRS; S.L. is Research Associate of the F.R.S-FNRS. The work performed in the Nguyen laboratory is supported by ULiège (Crédit Classique), the F.R.S.-FNRS (PDR T.0185.20; EOS 0019118F-RG36), the WEL Research Institute (CR-2022A-12), the Fonds Leon Fredericq, the Fondation Simone et Pierre Clerdent, the Fondation Médicale Reine Elisabeth, the ERANET Neuron (STEM-MCD and NeuroTalk), the Win2Wal (ChipOmics; #2010126), and the ERC-Synergy (UNFOLD).

## Supplemental Figure legends

**Figure S1.**
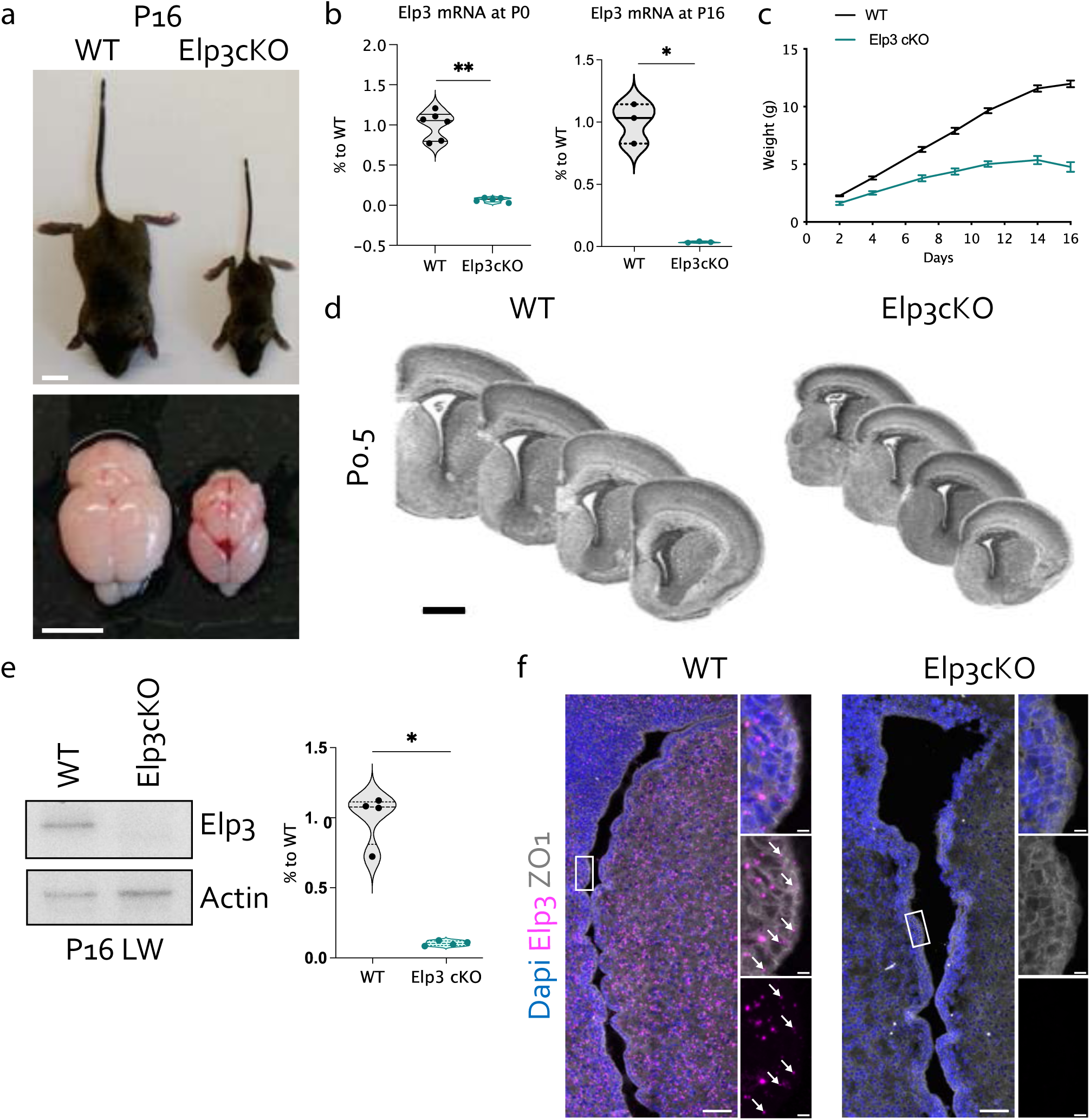
(links to Figure 1). Characterization of Elp3cKO mouse brain **a**, P16 old Elp3 cKO mice are undersized compared to littermate controls. They have a smaller brain with proportionally more reduction of its telencephalic vesicles. Scale bars, above: 1 cm; under: 5mm. **b,** Violin plot of Elp3 mRNA expression at P0 with median ± quartiles from six WT and five Elp3cKO mice. p-values were determined by Mann-Whitney test p=0.0043. Violin plot of Elp3 mRNA expression at P16 with median ± quartiles from three WT and Elp3cKO mice. p-values were determined by Mann-Whitney test p=0.05. **c,** Weight curve of WT and Elp3cKO mice from P2 to P16, mean ± s.e.m from four WT and three Elp3cKO mice. **d**, Consecutive rostro-caudal coronal sections of P0 WT and Elp3cKO mouse telencephalon. Scale bars, 500µm. **e,** Western blotting on Elp3 WT and Elp3 cKO P16 LW extracts. Violin plots show median ± quartiles from five (WT) and four (Elp3cKO) biological replicates. p-values were determined by Mann-Whitney test \**p*[=[0.0286. **f,** Confocal pictures of WT and Elp3cKO P7 lateral wall. High power magnification shows Elp3 mRNA expression that are lost in Elp3cKO. Scale bars, 50µm and 5µm on the zoom.

**Figure S2.**
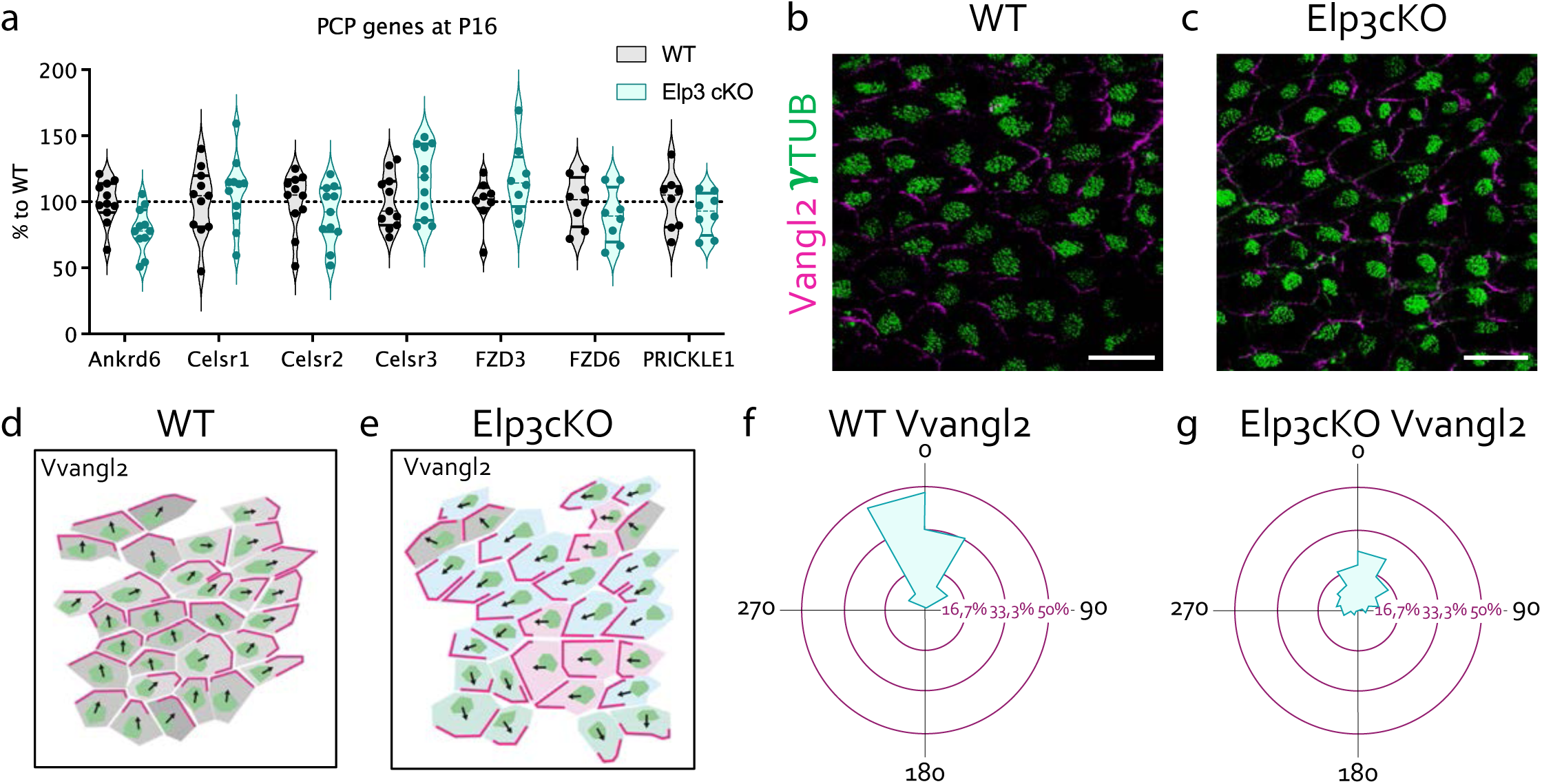
(links to Figure 2). Loss of Elp3 affects tissue level distribution of planar cell polarity proteins **a,** Violin plot of the mRNA expression of the main gene effectors of planar cell polarity at P16 with median ± quartiles from eight to eleven WT and Elp3cKO mice. P-values were determined by two-way ANOVA with Sidak’s multiple comparisons test, no significant results were observed. **b,c,** Immunolabeling of cilia patches detecting γ-tubulin (green) and the PCP protein Vangl2 (magenta) on WT (**b**) and Elp3cKO (**c**) brain lateral wall of P16 mice. Scale bars, 20µm. **d,e,** A vector (Vvangl2) pointing from the geometrical center of each cilia patch (green) to the middle of the Vangl2 staining (magenta) was drawn, and can be found opposite of each other according the immunolabeling for γ-tubulin and Vangl2 in P16 WT (**d**) and Elp3cKO mice (**e**). **f, g,** Circular dispersion of Vvangl2 around the mean in WT (288 cells) and Elp3cKO (587 cells) brain lateral wall from three mice of each genotype. p values were determined by Watson U^2^ test=4.113, *** p<0.001.

**Figure S3.**
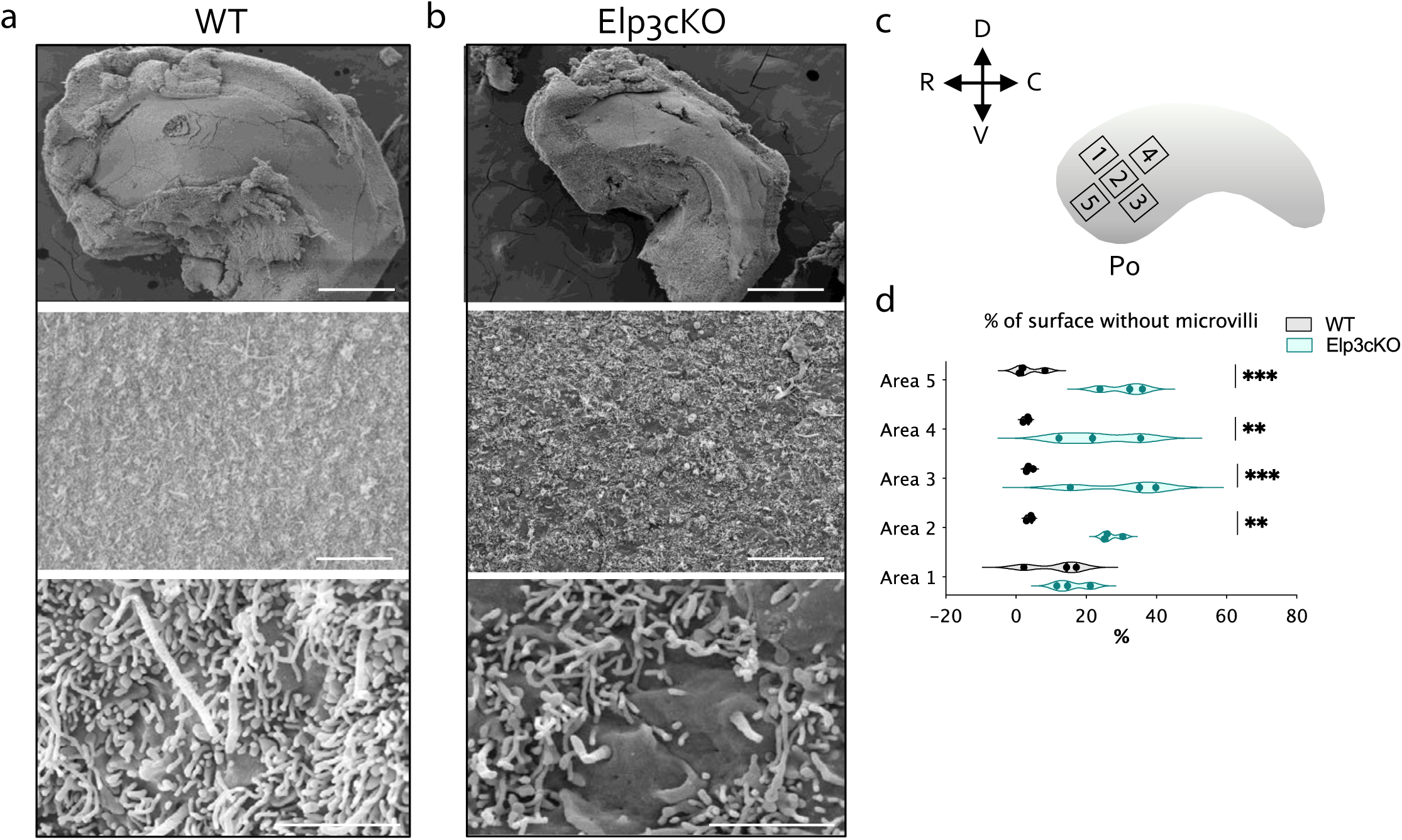
(links to Figure 3). The apical surface of maturating ependymal cells show reduced microvilli number upon loss of Elp3 expression **a,b,** Scanning electron microscopy micrograph of P0 WT (**a**) and Elp3cKO (**b**) brain lateral walls. Progressive close up are provided from top to bottom. Scale bars top, 500µm; Scale bars middle, 10µm and Scale bars bottom 2µm. **c,** Schematic representation of the ventricular lateral wall, depicting the different quantified areas for the microvilli quantification of the scanning electron microscopy images. **d,** Violin plot shows the microvilli coverage at P0 with median ± quartiles from three WT and Elp3cKO biological replicates. P-values were determined by two-way ANOVA with Sidak’s multiple comparisons test ** p<0.01 and *** p<0.001.

**Figure S4.**
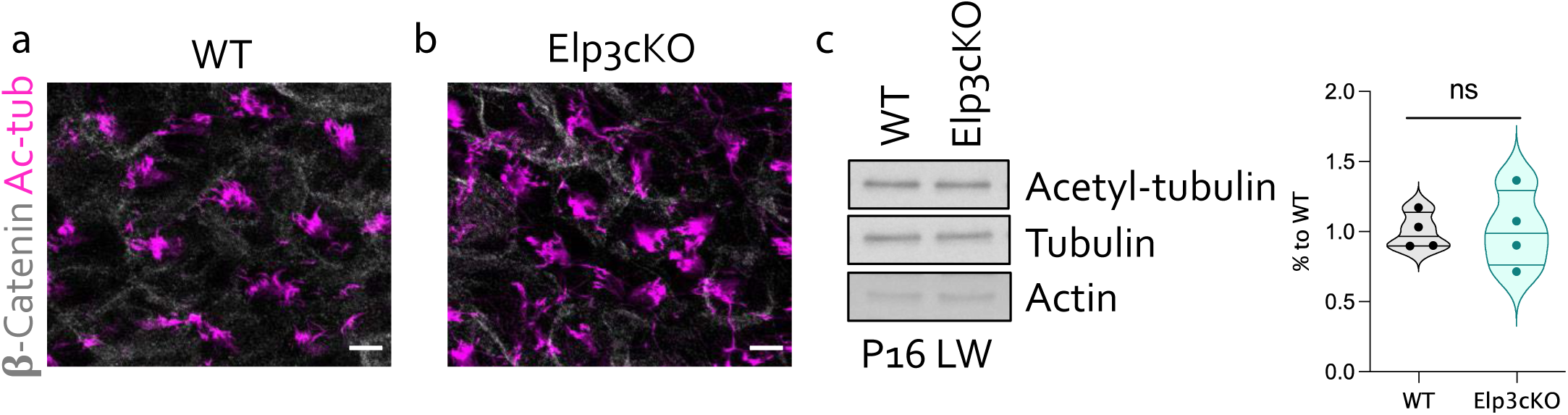
(links to Figure 4). Loss of Elp3 does not alter tubulin acetylation in brain lateral wall extracts **a,b,** Immunolableling of ependymal cells showing motile cilia labelled expressing acetyl-tubulin (magenta) and cell borders delineated by β-catenin (light grey) on P16 WT (**a**) and Elp3cKO (**b**) brain lateral walls. Scale bars, 10µm. **c,** Western blotting of acetyl-tubulin, tubulin and actin on P16 WT and Elp3cKO lateral wall extracts. Violin plots of acetyl-tubulin/tublin ratios, median ± quartiles from four mice of each genotype, as reported on the figure. P values were determined by Mann Whitney test, no significant difference was observed.

**Figure S5.**
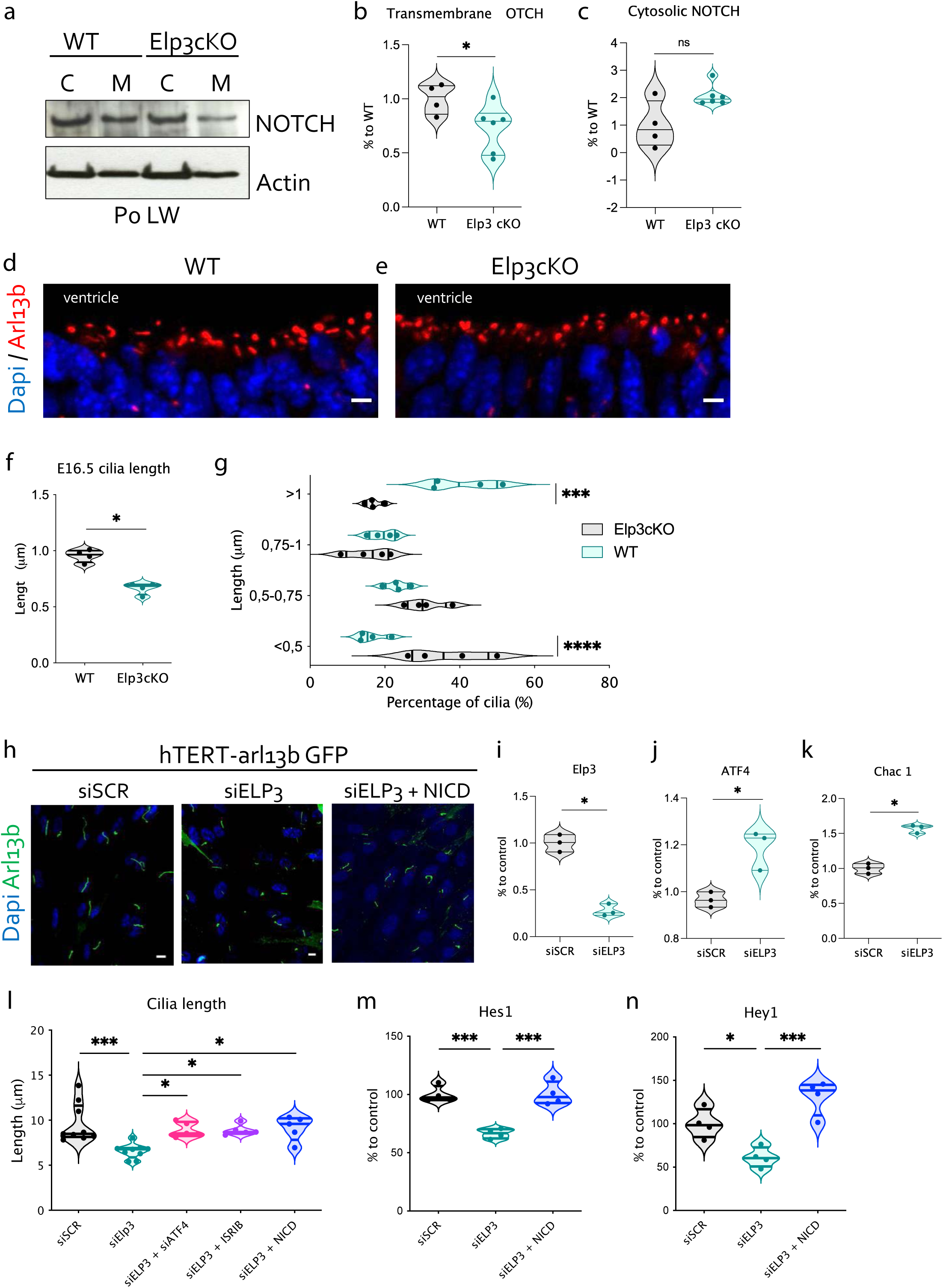
(links to Figure 4). Loss of Elp3 triggers ER/UPR and interferes with Notch signalling in ciliated cells **a,** Western blotting of NOTCH1 and actin on P16 WT and Elp3cKO cytosolic and transmembrane fraction extracts from lateral wall extracts. **b,c,** Violin plots of the expression of transmembrane and cytosolic NOTCH1, respectively with median ± quartiles from four WT and six Elp3cKO mice. P values were determined by Mann Whitney test, * p=0.0381. **d,e,** Immunolabeling of primary cilia cells labelled with arl13b (red) and dapi (blue) on E16.5 WT (**d**) and Elp3cKO (**e**) embryo. Scale bars, 5µm. **f,** Primary cilium length at E16.5 for WT and Elp3cKO. Violin plots of RGC primary cilium length, median ± quartiles from four mice for each genotype. p values were determined by Mann Whitney test p=0.0286. **g,** Primary cilium length quantiles at E16.5 for WT and Elp3cKO mice. Violin plots of primary cilium length, median ± quartiles from four mice for each genotype. p values were determined by two-way ANOVA with Sidak’s multiple comparisons test \*\*\**P*<0.001 and \*\*\*\**P*<0.0001. **h,** Human hTERT-RPE1 cells that overexpress GFP-tagged Arl13B protein to visualize primary cilia transfected with esiRNA Elp3 (siElp3) with or without NICD overexpression compared to esiRNA control (siSCR) **i-k,** Violin plots of mRNA expression levels for Elp3 (**i**), ATF4 (**j**) and Chac1 (**k**), respectively in hTERT-RPE1 transfected with siSCR or siElp3. All violin plots show median ± quartiles for three biological replicates for each transfection conditions. All p values were determined by Mann-Whitney test p=0.05. **l,** Primary Cilium length in hTERT-RPE1 tranfected with siSCR, siElp3, siElp3 combined with either siATF4, with/without ISRIB treatment or with/without NICD expression. Violin plots show median ± quartiles from five to nine biological replicates. p values were determined by One-way anova with Tukey’s multiple comparisons test ***p<0.001 and *p<0.05. **m-n,** Violin plots of mRNA expression levels for NICD responsive genes Hes1 (**m**) and Hey1 (**n**), respectively in hTERT-RPE1 transfected with siSCR, siElp3 and siEpl3 with/without NICD expression. All violin plots show median ± quartiles for four biological replicates. p values were determined by One-way anova with Tukey’s multiple comparisons test \*\*\**p*<0.001.

**Figure S6.**
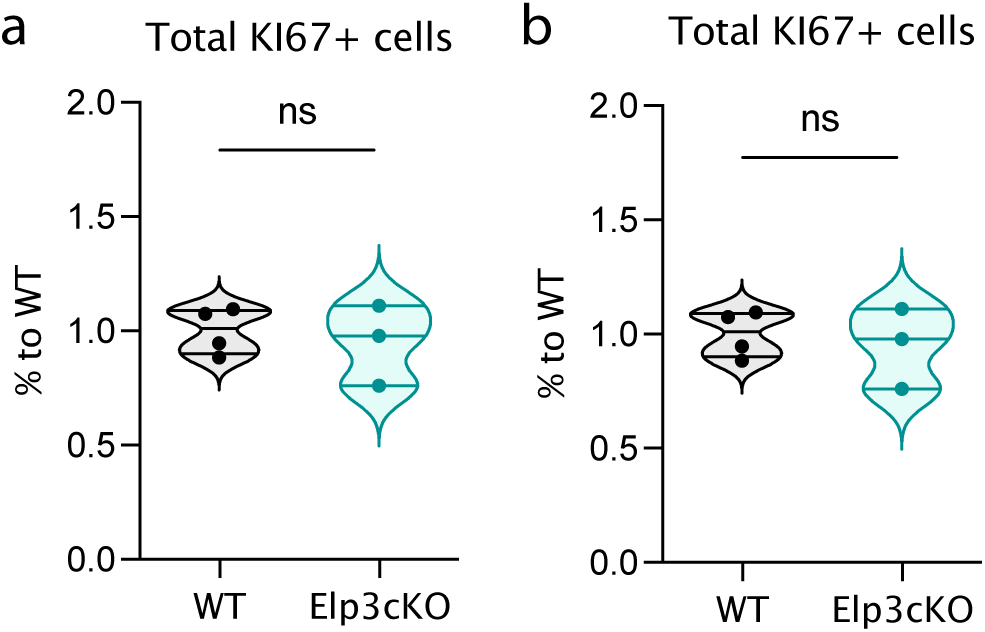
(links to Figure 5). The percentage of cycling of RGCs is not affected in mouse embryo upon loss of Elp3 **a,b,** Violin plots of EdU+cells and KI67+ RGC cells, respectively of E16.5 WT and Elp3cKO embryos fixed 24 hours after the EdU pulse. Violin plots show median ± quartiles from four WT and three Elp3cKO mice per genotype. p values were determined by Mann-Whitney test, no significant difference was observed.

